# CD29 marks superior cytotoxic human CD8^+^ T cells

**DOI:** 10.1101/562512

**Authors:** Benoît P. Nicolet, Aurélie Guislain, Floris P.J. van Alphen, Raquel Gomez-Eerland, Ton N. M. Schumacher, Maartje van den Biggelaar, Monika C. Wolkers

## Abstract

Cytotoxic CD8^+^ T cells can effectively kill target cells by producing cytokines, chemokines and granzymes. Expression of these effector molecules is however highly divergent, and tools that identify and pre-select potent killer cells are lacking. Human CD8^+^ T cells can be divided into IFN-γ and IL-2 producing cells. Unbiased transcriptomics and proteomics analysis on cytokine-producing fixed CD8^+^ T cells revealed that IL-2^+^ cells produce helper cytokines, and that IFN-γ^+^ cells produce cytotoxic molecules. IFN-γ^+^ T cells could be identified with the surface marker CD29 already prior to stimulation. CD29 also marked T cells with cytotoxic gene expression from different tissues in single-cell RNA-sequencing data. Notably, the cytotoxic features of CD29^+^ T cells were maintained during cell culture, suggesting a stable phenotype. Pre-selecting CD29-expressing MART1 TCR-engineered T cells potentiated the killing of target cells. We therefore propose that CD29 expression can help evaluate and select for potent therapeutic T cell products.

## INTRODUCTION

CD8^+^ T cells can effectively clear cells from our body that are infected with intracellular pathogens. In addition, CD8^+^ T cells are critical for immunosurveillance to eliminate pre-cancerous or cancerous cells. To exert their effector function, T cells produce pro-inflammatory molecules such as IFN-γ and TNF-α, granzymes and perforin, and chemokines such as CCL3 (MIP1α) and CCL4 (MIP1β) (Donia et al., 2017; Xue et al., 2017; Ma et al., 2013). Intriguingly, human T cells do not respond uniformly to activation, but rather show a heterogeneous production profile of effector molecules (Han et al., 2012; Nicolet et al., 2017). This may derive from stochastic and oscillating production of effector molecules, as suggested in developmental processes (Huang, 2009). However, it may also be due to intrinsic differences in the effector response to antigen by distinct CD8^+^ T cell types.

Interestingly, even though activated human CD8^+^ T cells efficiently produce IFN-γ and IL-2, we and others found that they generally produce only one of the two cytokines (Frentsch et al., 2013; Nicolet et al., 2017; Hamann et al., 1997). Only a small proportion of human CD8^+^ T cells produces IFN-γ and IL-2 combined (Frentsch et al., 2013; Nicolet et al., 2017; Hamann et al., 1997). Several studies proposed that human and mouse CD8^+^ T cells can provide T cell ‘help’ by expressing CD154/CD40L (Frentsch et al., 2013; Wong et al., 2008; Hernandez et al., 2007), in addition to their well-known cytotoxic function. CD40L-expressing human CD8^+^ T cells were shown to produce IL-2, but not IFN-γ (Frentsch et al., 2013). However, whether this cytokine production profile is stochastic, or whether IFN-γ and IL-2 producing CD8^+^ T cells represent two different T cell subsets and/or a divergent differentiation status is yet to be determined. It is also not known whether IL-2 and IFN-γ producing CD8^+^ T cells possess a different cytotoxic potential.

In this study, we set out to dissect the properties of human CD8^+^ T cells that display a differential cytokine production profile. Previous studies used cytokine capture assays to isolate T cells for gene expression analysis (Okoye et al., 2014). While this method is very accurate for low abundance responses, the risk of selecting false-positive cells by capturing cytokines from the neighbouring cell is substantial. We therefore developed a protocol that allowed for efficient isolation of RNA and protein from fixed T cells after intracellular cytokine staining. With this top-down approach, we performed an unbiased RNA-seq and Mass spectrometry analyses on IFN-γ and IL-2 producing primary human CD8^+^ T cells.

We found that IFN-γ producing T cells exhibit a cytotoxic expression profile, which is sustained during T cell culture. The surface molecules CD29 and CD38 identified the cytotoxic IFN-γ-producing T cells, and the non-cytotoxic IL-2 producing CD8^+^ T cells, respectively, a feature that is maintained during prolonged T cell culture. Using these two surface markers allowed us to distinguish T cells with a distinct cytokine production profile and cytotoxic potential already prior to T cell stimulation. Furthermore, the CD29 gene expression signature was a good prognostic marker for long term survival in melanoma patients. In conclusion, we here provide evidence that CD29 selects for bona fide cytotoxic CD8^+^ T cells, a marker that could potentially be used to evaluate the quality of therapeutic T cell products and possibly to select for effective adoptive T cell products.

## RESULTS

### Efficient recovery of RNA and protein from T cells after intracellular cytokine staining

To determine whether and how the gene expression profile of IFN-γ producing CD8^+^ T cells differed from IL-2 producing cells, we needed a method to separate high percentages of cytokine producers while preventing possible contaminations from false-positive cells. We therefore developed a protocol for efficient recovery of RNA and proteins from FACS-sorted fixed cells after intracellular cytokine staining. With the standard TRIZOL RNA isolation protocol, >99% of the mRNA isolated from PMA-ionomycin activated and formaldehyde-fixed T cells was lost, as determined by RT-PCR of several standard housekeeping genes, and of *IFNG* and *IL2* mRNA (Fig. S1A, B). To conserve the RNA integrity, we performed the intracellular cytokine staining (ICCS) and FACS in an RNA-protecting high salt buffer that contained RNAse inhibitors (Fig. 1A). This alteration in the protocol preserved the quality of ICCS, and thus the distinction of different cytokine producers (Nicolet et al., 2017). To free RNA from protein complexes, we included a proteinase K digestion step prior to RNA isolation (Nilsson et al., 2014) (Fig. 1A). These adjustments yielded mRNA levels from fixed T cells comparable to “fresh” T cells that were prepared without permeabilization and formaldehyde fixation (Fig. S1A, B). Importantly, the RNA purity and integrity was compatible with RNA-seq analysis, as revealed by nanodrop analysis and RNAnano Chip assay (Fig. S1C).

**Figure 1:**
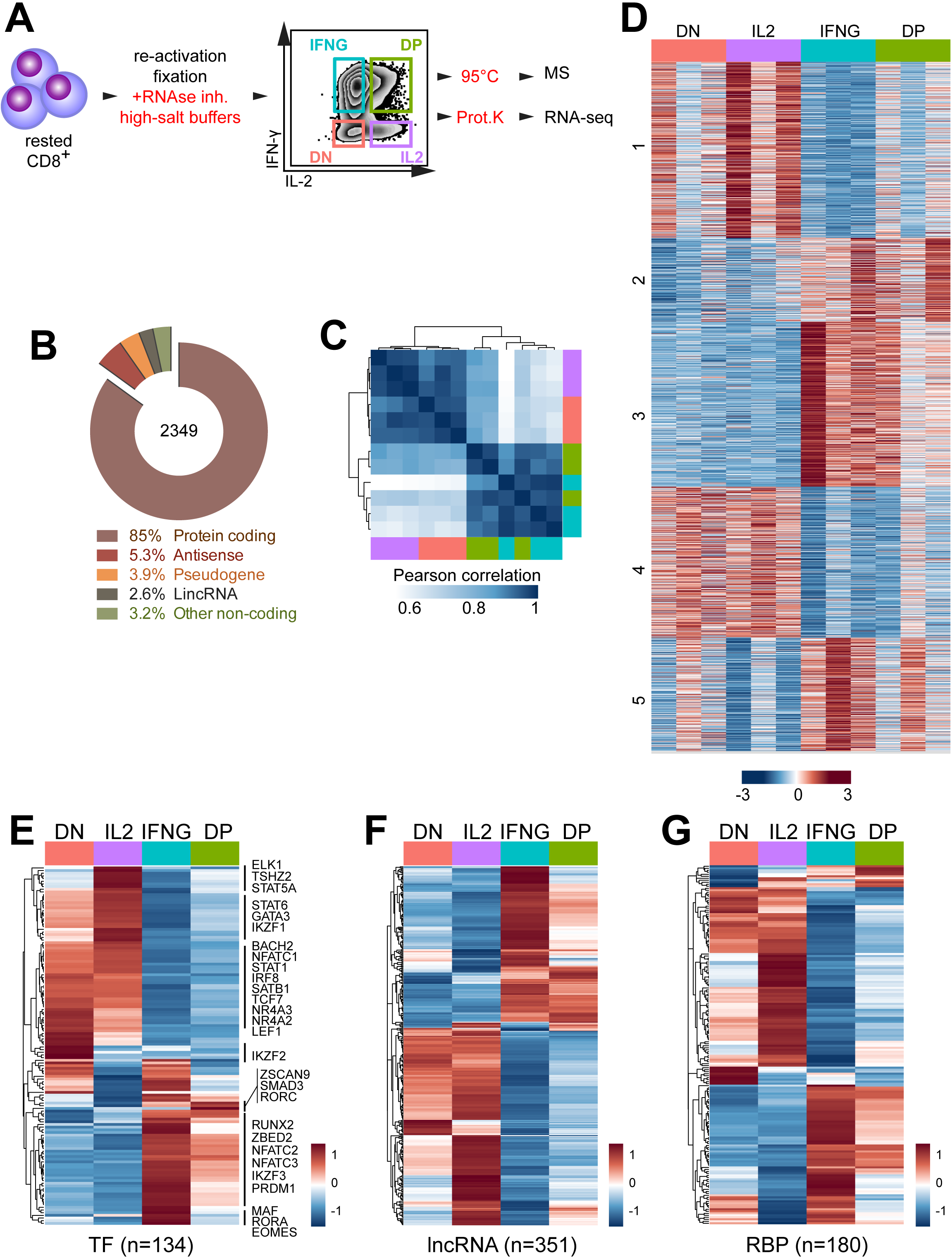
RNA-seq analysis on CD8^+^ T cells after intracellular cytokine staining. (**A**) Scheme to isolate RNA and protein from cytokine-producing, fixed T cells. Primary human CD8^+^ T cells were activated for 48h with αCD3/αCD28, and rested for 4 days in the presence of 10ng/mL rhIL-15. T cells were reactivated for 4h with 10ng/mL PMA and 1µM Ionomycin. Intracellular cytokine staining was performed in RNA-protecting buffers. Cells were FACS-sorted under RNA-protecting conditions. Total RNA and protein was recovered for RNA-seq and Mass Spectrometry analysis (Fig. 2) by reversing formaldehyde cross-linking. *IL2*: IL-2 single producers; *IFNG*: IFN-γ single producers; *DP*: double positive: IL-2 and IFN-γ co-producers; *DN*: double negative for IL-2 and IFN-γ production. (**B-H**) RNA-seq analysis of CD8^+^ T cells with a differential IL2 and/or IFN-γ production profile. (**B**) Gene biotypes (n=2349) that are differentially expressed between all 4 T cell populations. (**C**) Sample-wise Pearson correlation coefficient of differentially expressed genes (n=2349). (**D**) Heatmap of differentially expressed protein-coding genes (n=1984), numbers indicate k-means clusters (k=5). (**E**) Heatmap of differentially expressed transcription factors (TF) with >100 reads per million (rpm; n=134), (**F**) long non-coding RNAs (lncRNA) (n=351), and (**G**) RNA-binding proteins (RBP) with >10rpm (n=180). (E-G) Expression levels of biological replicates were averaged (n=3 per population).

To achieve mass spectrometry (MS)-grade protein recovery, we used the Filter Aided Sample Preparation (FASP) protocol (Wiśniewski et al., 2009). This protocol is compatible with MS analysis from formalin fixed material, as it contains a 95°C heating step to reverse the formaldehyde cross-linking of the ICCS procedure (Shi et al., 2006; Magdeldin and Yamamoto, 2012). Indeed, the correlation coefficient of MS analysis from fixed and fresh samples was r=0.973 (Fig. S1D). Thus, RNA and protein could be efficiently recovered from cytokine-producing, formaldehyde-fixed T cells, which allowed us to study their expression profile in depth.

### Differential gene expression profile of IFN-γ and IL-2-producing CD8^+^ T cells

To determine the gene expression profile of IFN-γ producing cells with that of IL-2 producing cells, human CD8^+^ T cells from 3 biological replicates, each containing a pool of 40 healthy donors, were activated for 2 days with α-CD3/α-CD28. T cells were removed from the stimulus and cultured for 4 days in the presence of human recombinant IL-15. ICCS after 4h of activation with PMA-ionomycin identified four T cell populations: IFN-γ producers (IFNG), IL-2 producers (IL2), IFN-γ/IL-2-coproducing T cells (double positive, DP), and T cells that did not produce detectable levels of either cytokine (double negative, DN; Fig. 1A).

Of the RNA-sequencing data, on average 19.8 million reads (∼92%) of the total 21.4 million reads could be mapped to the genome of each T cell population. Of these, 2349 genes were differentially expressed in at least one population, with a cut-off of log2 fold change >0.5, p-adjusted <0.01 (table S1). According to Ensembl-BioMart gene annotation, 85% (n=1984) of the differentially expressed genes were protein coding, 5.3% were antisense, 3.9% were pseudogenes, 2.6% were long-intergenic non-coding RNA (lincRNA), and 3.2% comprised other classes of non-coding RNA (ncRNA; Fig. 1B).

Pearson’s correlation coefficient of differentially expressed genes revealed that each biological replicate clustered according to its cytokine production profile (Fig. 1C). Interestingly, IFNG cells clustered with DP cells, and IL2 cells with DN cells (Fig. 1C). This close relationship of IFNG and DP cells and of IL2 with DN cells was also revealed by gene cluster analysis using k-means (k=5), in particular in cluster 2 and 4 (241 and 434 genes, respectively) (Fig. 1D, table S1). Yet, cluster 1 (507 genes) was more specific for IL2 T cells, and for IFNG T cells cluster 3 and 5 (474 and 326 genes, respectively; Fig. 1D, table S1).

We next interrogated whether gene expression modulators like transcription factors (TFs), long non-coding RNAs (lncRNAs), and RNA binding proteins (RBPs) were differentially expressed in IFNG and IL2 producers. Most of the 134 differentially expressed TFs were shared between DN and IL2 producers, and between IFNG and DP producers (Fig. 1E, table S2). This included GATA3 and IRF8 for IL2/DN producers, and PRDM1 (Blimp-1), NFATc2 and NFATc3 for IFNG/DP producers (Fig. S1E, table S2). Specific gene expression was detected for Eomes and RORA in IFNG producers, and for RORC (ROR-γt) in DP producers (table S2). A small cluster of TFs including IKZF2 (Helios) was specifically up-regulated in DN cells (Fig. S1E). Also the 351 differentially expressed lncRNAs revealed the close kinship between DN/IL2 producers and IFNG/DP producers (Fig. 1F, table S2). For instance, the IFN-γ promoting IFNG-AS1 (NEST) (Gomez et al., 2013) was highly expressed in IFNG/DP producers (Fig. S1F). Conversely, GATA3-AS1 known to re-enforce the Th2 phenotype in CD4^+^ T cells (Gibbons et al., 2018) showed highest gene expression in IL2 producers (Fig. S1F).

RBPs are critical regulators of RNA splicing, stability and translational control (Newman et al., 2016). Again, most of the 180 differentially expressed RBPs were similarly expressed in IFNG/DP producers and in IL2/DN producers (Fig. 1G, table S2). Combined, these data revealed a differential gene expression profile in IFNG and IL2 producers, which correlates with distinct expression of gene regulators.

### Differential protein signature of IL-2 and IFN-γ producing CD8^+^ T cells

We next determined the protein expression profile of the four FACS-sorted CD8^+^ T cell populations by MS. We identified in total 5995 proteins from the 3 biological replicates. After removing possible contaminants and filtering for proteins that were present in each biological replicate in at least one of the four populations, we identified a total of 3833 proteins. Each sample contained >3500 identified proteins (Fig. 2A, table S1). Pearson’s correlation of the MS analysis also revealed the kinship between IFNG and DP producers, and between IL2 and DN producers (Fig. 2B), and by k-means clustering of the 81 differentially expressed proteins (9 clusters; Fig. 2C). Of note, 42 (52%) of the differentially expressed proteins overlapped with the differential gene expression profile, including IL-2 and IFN-γ (Fig. 2D). In conclusion, IL2 and IFNG producers display a distinct gene and protein expression signature.

**Figure 2:**
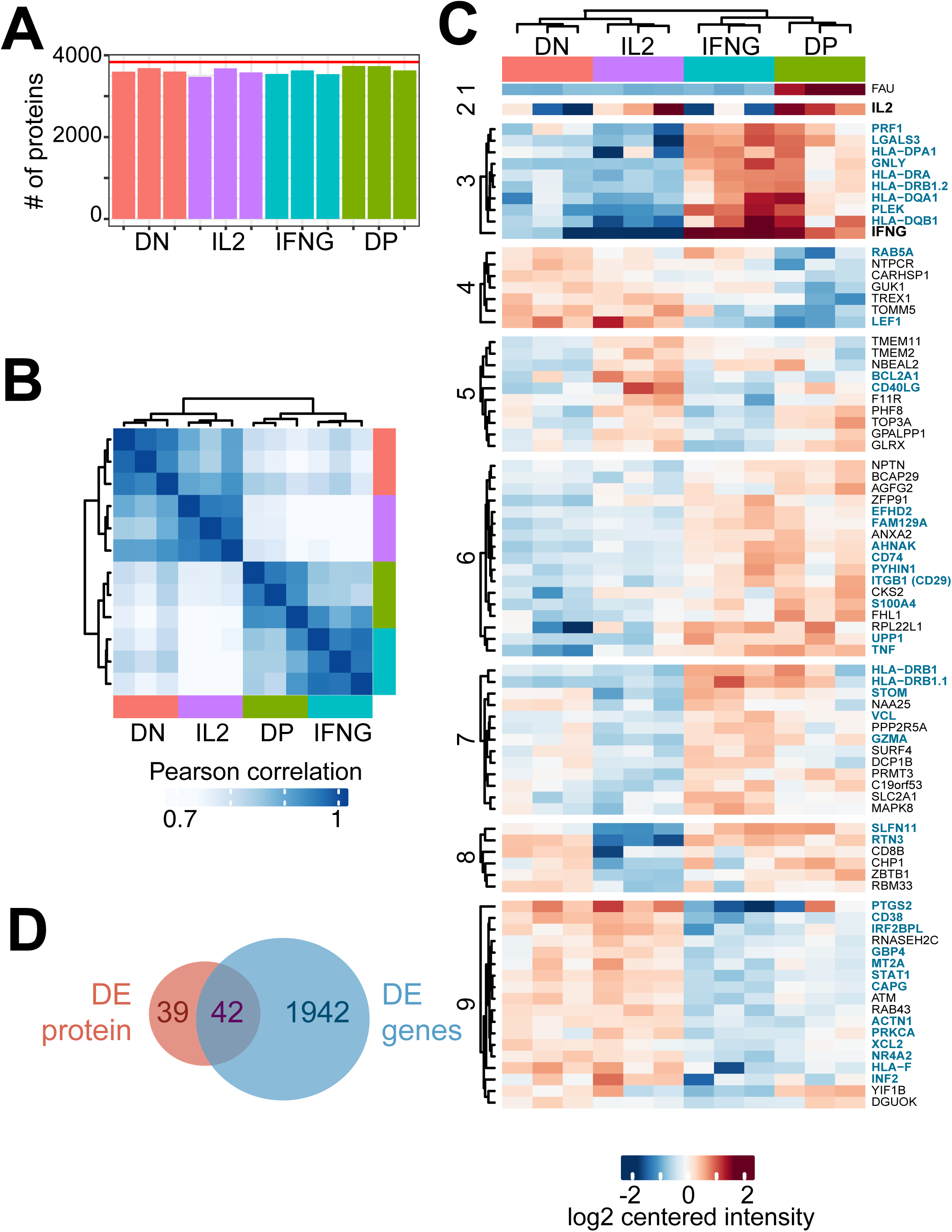
Mass-spectrometry analysis on cytokine producing, fixed CD8^+^ T cells. **(A)** Number of identified proteins by Mass-spectrometry analysis of FACS-sorted IL2, IFNG, DN and DP producers per sample (bars) and of all samples combined after filtering (n=3833, red line). **(B)** Sample-wise Pearson correlation coefficient of (n=81) differentially expressed proteins. Color-coding of cell populations as in (A). (**C**) Heatmap of differentially expressed proteins, numbers indicate k-means clusters (k=9). (**D**) Venn diagram of differentially expressed proteins and genes (fig.1). Proteins that show differential expression in mRNA and protein are indicated in (D) in blue.

### IFN-γ producing CD8^+^ T cells display a cytotoxic profile

To determine whether IFNG and IL2 producers showed a distinct gene expression also of other secreted effector molecules, we selected differentially expressed genes from Fig. 1 that 1) had expression levels >100 RPM and that 2) were annotated as “secreted” in the human protein atlas (Uhlén et al., 2015). After removing nuclear components and mitochondrial and ribosomal proteins (see method section), we identified 67 differentially expressed genes encoding secreted proteins (Fig. 3A). For IL2 producers, these included the prototypic Th2 helper effector molecules IL-4, IL-3, leukemia inhibitory factor (LIF) (Mosmann et al., 1986; Piccinni et al., 1998), and the Th2-inducing cytokine IL-11 and prostaglandin-endoperoxide synthase 2 (PTGS2, or COX-2) (Iñiguez et al., 1999; Curti et al., 2001) (Fig. 3A). IL-2 and PTGS2 were also found differentially expressed in the MS analysis (Fig. 3B). Conversely, IFNG producers - and in most cases also DP producers - contained substantially higher gene expression levels of the prototypic cytotoxic molecules TNF-α, Granulysin, Granzyme A, Granzyme H and Perforin (Fig. 3A). Also the chemokines CCL5 (RANTES), CCL3 (MIP1α), CCL4 (MIP1β) and its alternative splicing variant CCL4L2, all of which are associated with antiviral and antibacterial activity of cytotoxic CD8^+^ T cells (Cocchi et al., 1995; Auerbach et al., 2012; Narni-Mancinelli et al., 2007), were highly expressed in IFNG and DP producers (Fig. 3A). IFN-γ, TNF-α, Granulysin, Perforin, and CCL5 were also identified by MS analysis (Fig. 3B).

**Figure 3:**
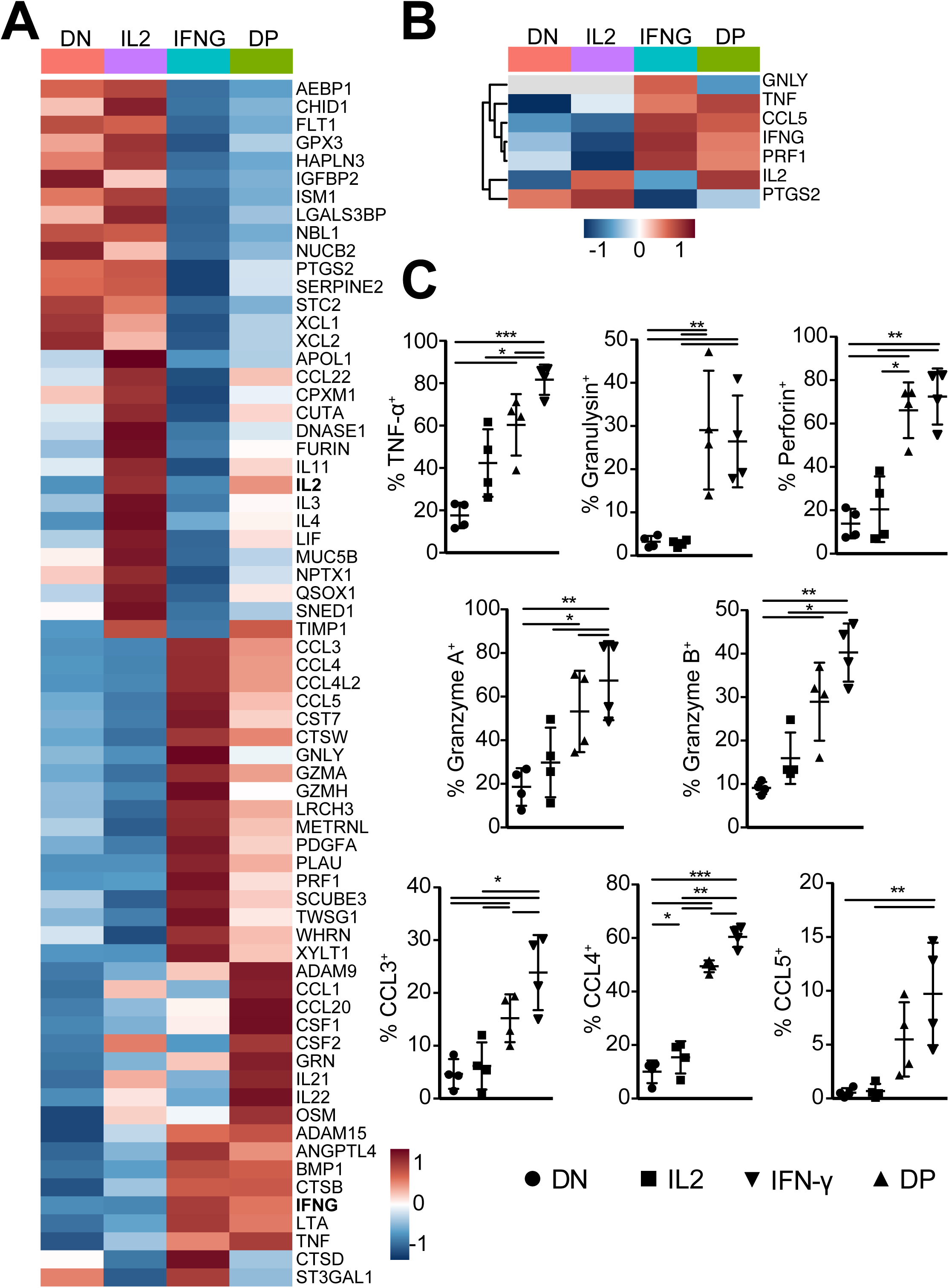
IL2 and IFNG producers express unique sets of secreted proteins. (**A-B**) Heatmap of differentially expressed genes (A, n=67) and proteins (B, n=7) that are annotated as secreted proteins (see mat/methods section). Expression levels of biological replicates were averaged (n=3 per population). (**C**) The production of secreted effector molecules identified in (A, B) was measured by intracellular cytokine staining in 4 new individual donors. Cells were activated with αCD3/αCD28 and rested as in Fig. 1, and reactivated with PMA-ionomycin for 4h in the presence of monensin to measure cytokine production. The percentage of indicated effector protein-producing cells in the IL2, IFNG, DP, and DN population was determined by flow cytometry. Repeated measurement ANOVA with Tukey post-test was performed. (*p<0.05, ** p<0.01, *** p<0.001).

We validated this prototypic cytotoxic gene expression profile of IFNG and DP cells in a new cohort of four individual donors. ICCS after reactivation for 4h with PMA-ionomycin confirmed that IFNG/DP producers produced significantly more TNF-α, Granulysin, Perforin, Granzyme A and B, and the chemokines CCL3, CCL4, and CCL5 than the DN/IL2 producers (Fig. 3C). Not only the percentage, but also the production of cytokines/chemokine per cell was higher, as determined by the geometric mean fluorescence intensity (geo-MFI; Fig. S2A). We therefore conclude that IFNG and DP CD8^+^ T cells preferentially express cytotoxic effector molecules.

### CD29 and CD38 expression preselects for IFN-γ and IL-2 producing CD8^+^ T cells

We next interrogated if surface markers could be used to identify IFNG producers and IL2 producers prior to T cell activation. To this end, we filtered the MS data for surface molecules that were ≥1.5 fold higher on average in expression in one of the four T cell populations. 19 surface markers showed differential protein expression in at least one of the four T cell populations (Fig. 4A). RNA-seq analysis identified 72 differentially expressed annotated proteins (Fig. S2B). We tested the expression pattern of 28 of these markers by flow cytometry on PMA-ionomycin activated CD8^+^ T cells on a new cohort of blood donors.

**Figure 4:**
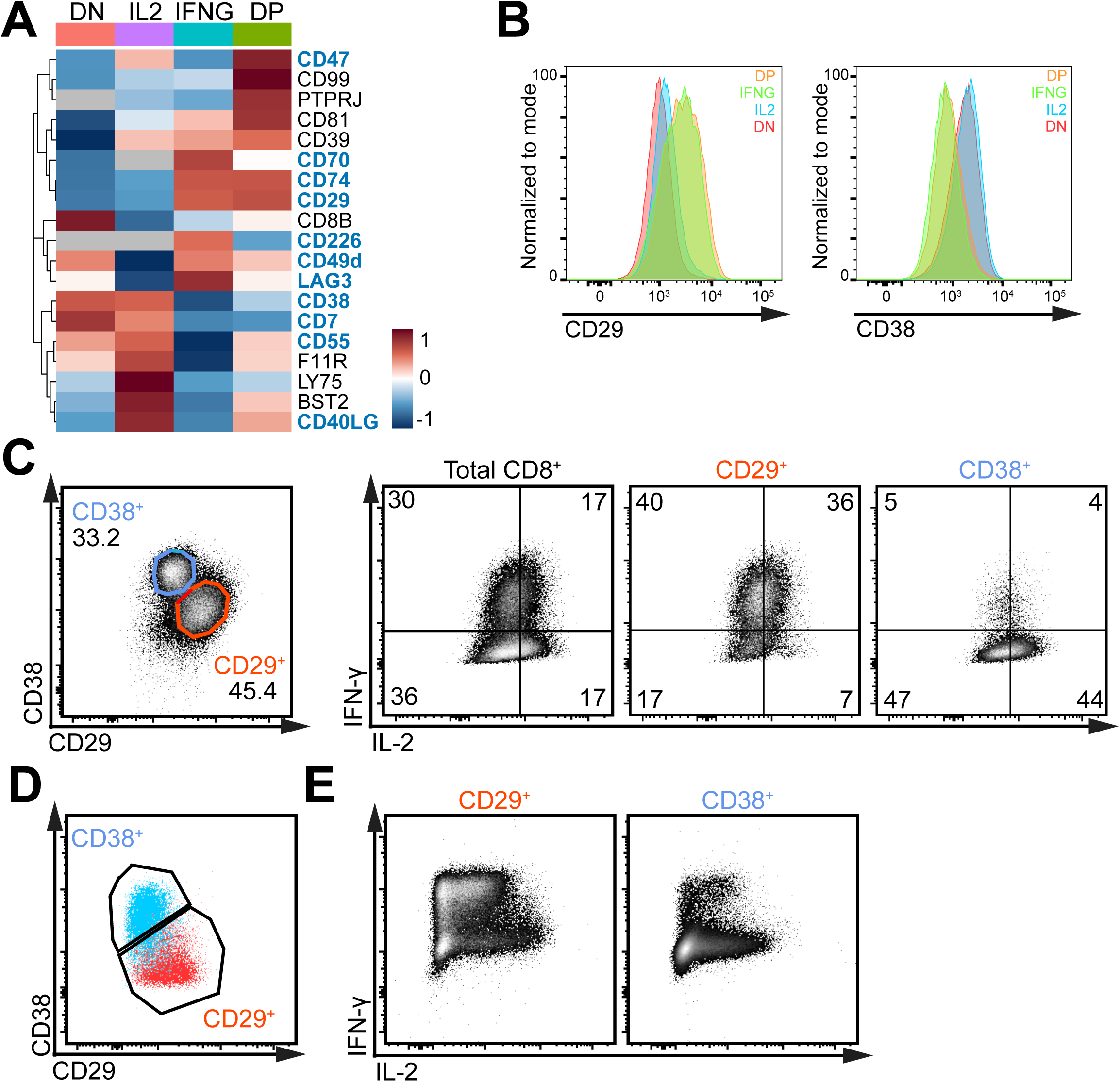
CD29 and CD38 identify IFNG and IL2 producers. **(A)** Heat map of differentially expressed CD molecules identified by MS analysis with LFC>1.5 (grey: not detected). Expression levels of biological replicates were averaged (n=3 per population). **(B)** CD29 (left) and CD38 (right) expression of CD8^+^ DN, IL2, IFNG and DP producers, as determined by flow cytometry after 4h reactivation with PMA-Ionomycin (n=6 donors). (**C**) Left panel: representative CD29 and CD38 expression of CD8^+^ T cells re-stimulated for 4h with PMA-Ionomycin. Right panel: IL-2 and IFN-γ production of total CD8^+^ T cell population, and of T cells gated based on CD29 and CD38 expression (representative of 17 donors). (**D-E**) Rested CD8^+^ T cells were FACS-sorted based on CD29 and CD38 expression and re-activated for 2 days with αCD3/αCD28, harvested and rested for another 21 days with rhIL-15. (**D**) Representative CD29 and CD38 expression at day 23 post sort. (**E**) Representative dot plot of IFN-and IL-2 production by CD29^+^ and CD38^+^ FACS-sorted CD8^+^ T cells that were reactivated for 4h with PMA-Ionomycin at day 23 post sort (representative of 4 donors).

None of the 16 surface markers that were only identified by RNA-seq analysis was suitable to select for IFNG and IL2 producers by flow cytometry (Fig. S2C). In contrast, the HLA class II histocompatibility antigen gamma chain CD74, and the β1-integrin were differentially detected in both MS and RNA-seq analysis (Fig. S2C) showed significantly higher cell surface expression in IFNG/DP producers than in IL2/DN producers (Fig. 4B; Fig. S2D, E). Conversely, the expression of CD40L, the complement decay-accelerating factor CD55, and the cyclic ADP ribose hydrolase CD38 were co-expressed and significantly higher in IL2/ DN producers (Fig. 4B, Fig. S2D, F).

In contrast to CD40L and CD74, CD29 and CD38 expression was also measured in non-activated CD8^+^ T cells. Importantly, the expression of CD29 and CD38 allowed for preselection of IFNG/DP producers and DN/IL2 producers even prior to activation (Fig. 4C). This included the production of TNF-α, Granzyme A, Granulysin, Perforin, CCL3, CCL4, and CCL5 by CD29^+^ T cells (Fig. S3C). Because CD29^+^ T cells included DP producers (Fig. 4B), the overall IL-2 protein production is similar in CD29^+^ and CD38^+^ T cells, yet clearly distributed between CD29^+^ DP and CD38^+^ IL2 single producers (Fig. S3D).

Intriguingly, even though the percentage of CD29^+^ and CD38^+^ T cells was greatly variable between donors (Fig. S3E), their expression pattern and cytokine production profile was maintained throughout time (Fig. 4D, E; Fig. S3F, G). Specifically, reactivation of T cells that were FACS sorted based on CD29 and CD38 expression on day 7 after activation and cultured for another 16 days showed a robust expression of CD29 and CD38, with CD29^+^ T cells being the prime IFNG producers, and CD38^+^ T cells the IL2 producers (Fig. 4D, E). CD29^+^ FACS-sorted T cells also expanded effectively in vitro (Fig. S3H). Using different cytokines during CD3/CD28 activation, or during the resting phase did not alter the percentage of CD29^+^ T cells (Fig. S3I, J). We therefore conclude that the differences between CD29^+^ and CD38^+^ T cells are cell-intrinsic and can be used to distinguish IFNG/DP producers from IL2/DN producers.

### CD29 is expressed on non-naïve CD8^+^ T cells

We next investigated whether CD29 expression also correlated with a cytotoxic gene signature of CD8^+^ T cells *ex vivo*. Because CD38 expression was not detected on peripheral blood-derived CD8^+^ T cells (Fig. 5A), we compared CD29^+^ with CD29^-^ T cells. Also *ex vivo*, CD29^+^ T cells were effective cytokine producers upon PMA-Ionomycin stimulation (Fig. S4A). In concordance with the described expression profile of cytotoxic molecules (Bengsch et al., 2018; Böttcher et al., 2015) and its expression in effector T cells (Sohen et al., 1990), CD29 expression was detected primarily in CD45RA^+^CD27^-^ Effector CD8^+^ T cells (T_Eff_), CD45RA^-^CD27^-^ Effector Memory (T_EM_) and CD45RA^-^CD27^+^ Memory CD8^+^ T cells (T_Mem_; Fig. 5B). Of the CD45RA^+^CD27^+^ naïve T cells (T_N_), only 6.6±4.2% expressed CD29 (Fig. 5B). However, further analysis revealed that the CD29^+^ ‘naïve’ T cells rapidly produce IFN-γ upon 4h PMA-Ionomycin stimulation, and express CD49d (Fig. 5C, D), previously shown to identify memory T cells with a naïve phenotype (TMNPs (Pulko et al., 2016)). This finding thus indicates that CD29 expression marks non-naïve CD8^+^ T cells.

**Figure 5:**
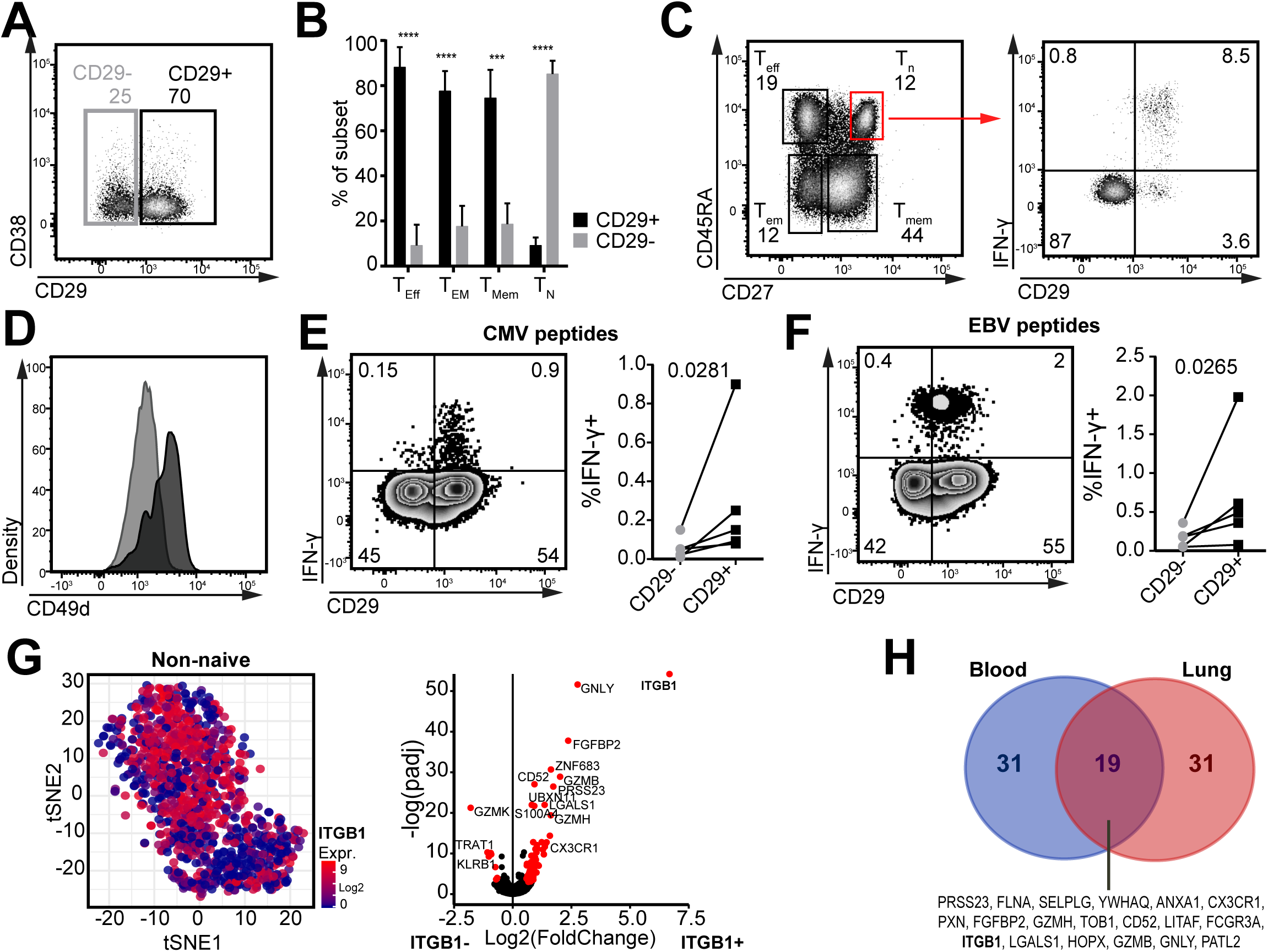
CD29 marks non-naïve CD8^+^ T cells ex vivo with a cytotoxic signature. (**A**) Representative expression profile of CD29 and CD38 of peripheral blood-derived CD8^+^ T cells *ex vivo*. (**B**) Percentage of CD29+ CD8^+^ T cell per subset, in blood, defined as: Naïve (CD27^+^CD45RA^+^), Memory (CD27^+^CD45RA^-^), Effector memory (CD27^-^CD45RA^-^) and Effector (CD27^-^CD45RA^+^, n=8 donors). (**C**) Representative dot plot of CD45RA and CD27 expression in CD8+ T cells activated with PMA-Ionomycin for 4h (left panel). T_N_ were gated and IFN-γ production was plotted against CD29 expression (right panel, representative of 5 donors). (**D**) *Ex vivo* CD49d expression of CD29+ (black) and CD29-(light grey) T_N_ CD8+ T cells from blood (representative of 4 donors). (**E**-**F**) Peripheral blood-derived CD8^+^ T cells were activated for 6h with MHC-I restricted peptide pools for CMV (**E**) or EBV (**F**). (E-F) representative dot plots of IFN-γ and CD29 expression (left panels) and quantification (right panels, n=5). (**G**) Single-cell gene expression analysis of peripheral blood-derived CD8^+^ T cells excluding naïve T cells (data retrieved from Single-cell RNA-seq data from(Guo et al., 2018), for details see Fig. S5B-F). Left panel: tSNE projection of ITGB1 (CD29) expression in CD8^+^ T cells excluding naïve T cells (n=1023, log2(TPM)). Right panel: volcano plot of differentially expressed genes in ITGB1^high^ and ITGB1^low^ CD8^+^ T cells. ITGB1 gene expression cut-off was determined as depicted in Fig. S4G. (**H**) Venn diagram of the core signature of CD29^+^CD8^+^ T cells from Blood and Lung (the top 50 up-regulated genes in each tissue). Differential expression with absolute(LFC)>0.5 and p-adjusted<0.05. (*p<0.05; **p<0.01).

To determine how CD29 expression related to antigen-specific triggering, we studied Choriomeningitis virus (CMV) and Epstein-Barr virus (EBV) specific T cells. In healthy blood donors, CMV-specific T cells are primarily of the effector, and EBV-specific T cells of the central memory phenotype (Appay et al., 2002; Hertoghs et al., 2010). We activated PBMCs with CMV and EBV peptide pools and found that the IFN-γ producing CMV- and EBV-specific T cells are primarily CD29^+^ (Fig. 5E, F). In conclusion, CD29 is almost exclusively expressed by non-naïve T cells.

### scRNA-seq analysis reveals the cytotoxic signature of CD29^+^ CD8^+^ T cells

To further characterize the gene expression profile of CD29^+^CD8^+^ T cells, we re-analysed single-cell RNA-seq (scRNA-seq) data of peripheral blood-derived CD8^+^ T cells from Guo *et al* (Guo et al., 2018). To distinguish naïve T cells from non-naïve CD29^+^ T cells, we used unsupervised clustering followed by differential expression analysis. We selected for high *LEF1*, *CCR7*, *SELL* gene expression and low *CCL5*, *GZMB*, *PRF1* gene expression to identify T cells with a T_N_ transcriptome signature (Willinger et al., 2006; Yang et al., 2016) (Fig. S4B, C). *CCL5* gene expression highly correlated with non-naïve T cells and was thus used to enrich for those for downstream analysis (Fig. S4D-F).

*ITGB1* (CD29) gene expression showed a non-uniform expression in non-naïve CD8^+^ T cells (Fig. 5G, left panel; Fig. S4G). We compared the gene expression profile of non-naïve T cells with high or low *ITGB1* gene expression, and found that high *ITGB1* gene expression strongly associated with high gene expression levels of *GNLY* (Granulysin), *GZMB* (Granzyme B), and the transcription factor *ZNF683* (Hobit) that is associated with cytotoxic activity of human T cells (Vieira Braga et al., 2015) (Fig. 5G, right panel). Conversely, in concordance with previously described gene expression profiles (Bengsch et al., 2018; Kragten et al., 2018), *ITGB1*^low^ CD8^+^ T cells associated with high gene expression levels of *GZMK* (Granzyme K), and the inhibitory receptor *KLRB1* (CD161; Fig. 5G, right panel, table S3).

scRNA-seq analysis of CD8^+^ T cells from lung tissue (Guo et al., 2018) also revealed a non-uniform expression of the *ITGB1* transcript (Fig. S4H, I left panel). Again, high *ITBG1* expression associated with a cytotoxic gene expression profile (Fig. S4I, right panel). When we compared the top 50 most upregulated genes in *ITGB1*^high^ T cells derived from blood and lung, and from lung and liver tissue (Zheng et al., 2017), we identified a core signature of *ITGB1*^high^ CD8^+^ T cells that encompassed the cytotoxic molecules *GNLY*, *GZMB*, *GZMH*, and *FGFBP2*, and *LGALS1* (Galectin-1; Fig. 5H, Fig. S4J). Of note, the fractalkine receptor *CX3CR1* is also part of the *ITGB1* ^high^ signature (Fig. 5H), which we confirmed with flow cytometry analysis of blood-derived CD8^+^ T cells (Fig. 5A). However, in contrast to murine CD8^+^ T cells (Böttcher et al., 2015), in humans CX3CR1 expression is almost exclusively found on CD45RA^+^CD27^-^ effector CD8^+^ T cells, and only on a minority of T_M_ or T_EM_ cells (Fig. S5B) (Nishimura et al., 2002). In line with the restricted expression pattern of CX3CR1, CD29 expression is superior in predicting the level of IFN-γ and TNF-α production (Fig. S5C-F). Thus, *ITGB1* gene expression identifies human T cells with a cytotoxic gene expression profile in blood and in tissues.

### CD29 gene signature is prognostic for long term survival of melanoma patients

We next interrogated if *ITGB1* gene expression correlated with good clinical responses. We focussed on cutaneous melanoma (SKCM), because the Cox regression analysis for survival from all TCGA datasets on CD8B expression identified SKCM as the only tumor type with a clear benefit of CD8^+^ T cell infiltration (Fig. S6A, p=0.000125). *ITGB1* is expressed by many cell types, including tumor cells (Sun et al., 2018). Therefore, we used the CD29^+^ core signature as defined in Figure 5H. Intriguingly, analysis of TCGA data of melanoma patients revealed that the CD29^+^ core signature correlated well with a positive clinical outcome (high CD29 signature: median=10.08 years, low CD29 signature: median= 4.84 years; p=<0.001; fig 6A). In particular, using the patient cohort with high CD8^+^ T cell infiltrates levels estimated by CIBERSORT (Newman et al., 2015) further showed a clear benefit of the CD29 core signature for long-term survival (high CD29 signature: median= 13.5 years, low CD29 signature: median= 5.37 years; Fig. 6B). Thus, the CD29 signature has a good prognostic value for melanoma patients.

**Figure 6:**
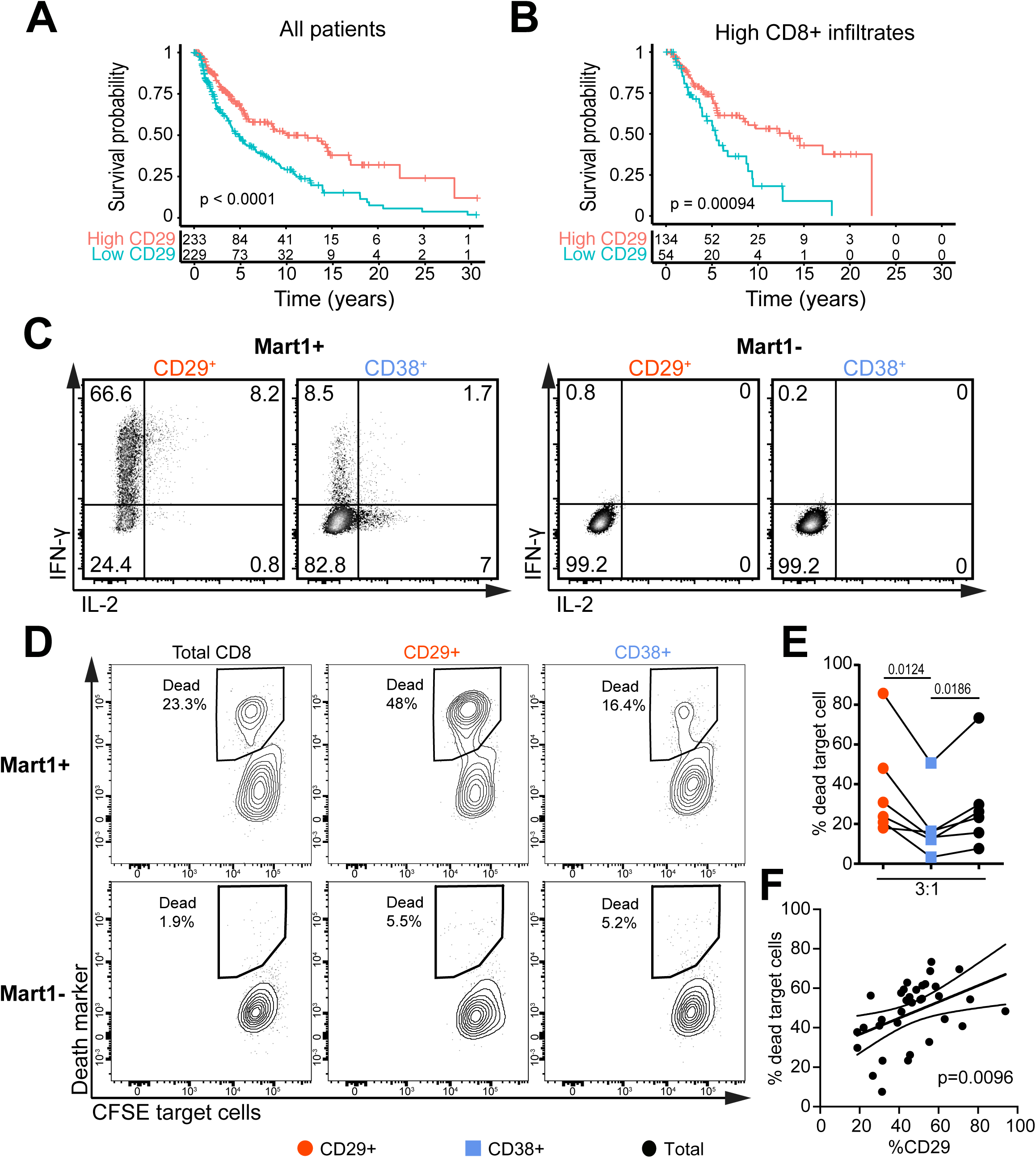
CD29 gene signature is associated with better survival, and CD29^+^T cells are superior in killing tumor cells. (**A-B**) Kaplan-Meier plot showing the overall survival of total skin cutaneous melanoma (SKCM) patients (**A**) or patients with high CD8^+^ T cell infiltrate estimates (**B**). Patients were stratified for a high or low CD29 gene signature as defined in Figure 5H (See method section). The lower panels depict the number of patient at risk for each time points. The indicated p-value was calculated using a log-rank test. (**C**) CD29^+^ or CD38^+^ MART1 TCR-engineered FACS-sorted CD8^+^ T cells were cultured for 6h with MART1+ or MART1-tumor cells at an effector: target (E:T) ratio of 3:1. Representative dot plot of IFN-γ and IL-2 production as measured by intracellular cytokine staining. (**D**) Total, or CD29^+^ and CD38^+^ FACS-sorted MART1-TCR expressing CD8^+^ T cells were co-cultured with CFSE-labeled MART1+ (upper panel) or MART1-(lower panel) tumor cells for 20h at a 3:1 E:T ratio. Representative measurements of dead tumor cells as determined by Near IR live-dead marker. (**E)** Tumor cell killing at 3:1 E:T ratio (n=6, repeated measurement ANOVA with Tukey post-test was performed). (**F**) Correlation of tumor killing of total CD8^+^ MART1-TCR expressing T cells with the percentage of CD29^+^CD8^+^ T cells present in the T cell product (n=34, linear regression).

### MART1 TCR-engineered CD29^+^ CD8^+^ T cells potently kill tumor cells

We then examined whether CD29 expression is indicative of effective T cell responses against tumor cells. We retrovirally transduced CD8^+^ T cells with the codon-optimized MART1-TCR that recognizes the HLA-A*0201 restricted 26-35 epitope of MART1(Gomez-Eerland et al., 2014). FACS-sorted CD29^+^ and CD38^+^ MART1 TCR-engineered CD8^+^ T cells were then exposed for 5h to a patient-derived HLA-A201^+^ MART1^high^ expressing melanoma tumor cell line (MART1+), or to the HLA-A201^-^ MART1^low^-expressing cell line (MART1-)(Marincola et al., 1994; Topalian et al., 1989) (Fig. 6C). MART1 TCR-engineered CD29^+^ T cells produced substantially higher levels of IFN-γ, TNF-α, Granulysin, and CD107α in response to MART1 antigen-expressing tumor cells than CD38^+^ T cells (Fig. S6B). CD29^+^ T cells were the major source of IL-2^+^IFN-γ^+^ double producing T cells, and CD38^+^ T cells were enriched for IL2 single producers (Fig. 6C, Fig. S6B).

Lastly, we determined whether MART1 TCR-engineered CD29^+^ T cells were also more potent in killing tumor cells than CD38^+^ T cells. CFSE-labelled MART1+ and MART1-tumor cells were co-cultured for 20h with FACS-sorted MART1 TCR-engineered total CD8^+^ T cells, or with FACS sorted CD29^+^ or CD38^+^ T cells. We found that MART1+ tumor cells, but not MART1-cells, were effectively killed by MART1 TCR-engineered T cells (Fig. 6D). Notably, in line with their cytotoxic gene and protein expression profile, CD29^+^ T cells showed superior killing when compared to CD38^+^ T cells (Fig. 6D, E). The superior killing was evident for different effector:target ratio’s (Fig. 6C).

Intriguingly, when we compared the killing capacity of the TCR engineered T cell product with the percentage of CD29-expressing cells present in the total T cell product, we found a clear correlation of these two parameters (Fig. 6F). Thus, CD29 expression identifies T cells with the highest cytotoxic potential in response to tumor cells.

## DISCUSSION

In this study, we present a method to perform RNA-seq and mass spectrometry analysis from formaldehyde-fixed, permeabilized primary CD8^+^ T cells. This method uses the endogenous protein expression measured by intracellular cytokine staining for selection, resulting in a reliable isolation of different populations without the need of genetic modifications (Okoye et al., 2014). We anticipate that one can readily adapt this rapid and cost-effective method to select cells with other features, such as differential chemokine or transcription factor expression.

Here, we uncovered that the differential cytokine production profile of human CD8^+^ T cells reflects stable T cell-intrinsic properties. Whereas IL2 producers display a gene expression profile resembling helper T cells, IFNG producers express a cytotoxic core signature. With CD29 and CD38, IFNG-DP producers can be identified and separated from DN-IL2 producers even prior to T cell stimulation, a feature that was maintained upon prolonged *in vitro* cultures. The division between highly cytotoxic CD29^+^ T cells and CD38^+^ T cells was not only observed in CD8^+^ T cells but was also true for CD4^+^ T cells (data not shown).

Importantly, scRNA-sequencing analysis on blood and lung derived CD8^+^ T cells revealed that the cytotoxic core signature of CD29^+^ T cells is not limited to *in vitro* T cell cultures but is also found *ex vivo*. It has been previously reported that a cytotoxic gene expression of tumor-infiltrating T cells is associated with better patient survival (Becht et al., 2016; Mlecnik et al., 2016). In line with that finding, TCGA analysis in cutaneous melanoma patients also indicates that the CD29 core gene signature is a good prognostic marker for long term survival. In fact, it further stratifies long-term survival in patients with high CD8 infiltration, which is already a good predictor.

We found that CD29 expression marks non-naive T cells. CX3CR1^+^ T cells are also included in this population. However, because CX3CR1 is primarily expressed in effector T cells in humans that are known to poorly expand in vitro (Böttcher et al., 2015), we consider CD29 superior to identify cytotoxic T cells with a good proliferative capacity. Selecting for CD29^+^ T cells resulted in the generation of TCR engineered T cell products with improved killing of target cells at least *in vitro*.

Between blood donors, we found a great variation in the percentage of CD29^+^ T cells. It is conceivable that this donor-individual difference in CD29 expression stems from a differential pathogenic and microbial exposure. Yet, CMV and EBV infections efficiently generated CD29^+^ IFN-γ-producing T cells, indicating that a variety of pathogens may yield cytotoxic CD29^+^ T cells. Of note, the clear correlation with CD29 expression and cytokine production was not detected in mice housed under specific pathogen free (SPF) conditions (unpublished observation). This may again point to the possibility that pathogen exposure contributes to the generation of non-naïve cytotoxic CD29^+^ T cells.

Previous studies showed that the cytokine production profile of CAR T cells was highly diverse (Xue et al., 2017). As culturing T cells with different cytokines did not alter the relative distribution of CD29^+^ and CD29^-^ T cells, we propose to select for CD29 expression for the generation of highly toxic T cell products, in particular when donors have a low percentage of CD29^+^ T cells. Selecting for CD29^+^ cytotoxic T cells to generate T cell products may also come with a positive side effect. IL2 producers displayed a Th2 gene expression profile, which includes IL-3, IL-4, IL-11, LIF and PTGS2 (COX-2). A Th2 cytokine profile may impede anti-tumoral responses (Chen and Mellman, 2013). In fact, IL2 and DN populations produce PTGS2 (COX-2) that was recently shown to suppress anti-tumoral immunity (Zelenay et al., 2015; Böttcher et al., 2018). It is therefore conceivable that removing IL2/DN producers from T cell products may already support the anti-tumoral responses of CD29^+^ T cells.

Another factor that impedes effective anti-tumoral T cell responses is the failure to migrate into the tumor tissue (Chen and Mellman, 2013; Mlecnik et al., 2016). Whereas β1 (CD29) and β2 integrins appear not to be required for T cells migration into at least some tumor types (Salmon et al., 2012), the expression of metalloproteinases is critical for this process (Ahrends et al., 2017). Intriguingly, we found that IFNG/DP producers also express higher transcript levels of the metalloproteinase ADAM15 and ADAM9, which could potentially support their intra-tumoral migratory capacity.

In summary, our data show that RNA-seq and Mass spectrometry analysis after intracellular cytokine staining is a powerful tool to define distinct T cell populations. This allowed us to uncover the differential make-up of CD8^+^ T cell responses that is conserved and sustained marked by CD29. Not only our findings may help improve T cell products for therapeutic purposes in the future, they provide a robust marker to study cytotoxic T cells.

## MATERIAL AND METHODS

### T cell activation

Peripheral blood mononuclear cells (PBMCs) from healthy volunteers were isolated by Lymphoprep (density gradient separation; StemCell) and stored in liquid nitrogen until further use. The study was performed according to the Declaration of Helsinki (seventh revision, 2013). Written informed consent was obtained (Sanquin, Amsterdam, The Netherlands).

For RNA-seq and MS analysis, cryopreserved blood from 3 independently obtained pools of 40 blood donors depleted for monocytes was used. CD8^+^ T cells were enriched to a purity of ±80% with the CD8 MACS positive isolation kit (Miltenyi). Validation essays were performed with cryopreserved blood from individual donors without MACS isolation.

T cells were activated as previously described (Nicolet et al., 2017). Briefly, non-tissue culture treated 24-well plates (Corning, USA) were pre-coated overnight at 4**°**C with 4μg/mL rat α-mouse IgG2a (MW1483, Sanquin) in PBS. Plates were washed and coated for >2h with 1μg/mL αCD3 (clone Hit3a, eBioscience) at 37**°**C. 0.8×10^6^ CD8^+^ enriched PBMCs/well were seeded with 1µg/mL soluble αCD28 (clone CD28.2, eBioscience) in 1mL IMDM (LONZA) supplemented with 8% Fetal Calf Serum, 100U/mL Penicillin, 100µg/mL streptomycin and 2mM L-glutamine. After 48h of incubation at 37**°**C, 5% CO_2_, cells were harvested, washed, and further cultured in standing T25/75 tissue culture flasks (Thermo-Scientific) at a density of 0.8×10^6^/mL, supplemented with 10ng/mL human recombinant IL-15 (Peprotech). Medium was refreshed every 5-7days. Alternatively, cells were cultured without cytokine or, with human recombinant IL-2 (50IU/mL), IL-6 (10ng/mL), IL-9 (10ng/mL), IL-11 (10ng/mL), IL-21 (50ng/mL).

### Preparation of activated and fixed T cells for RNA-seq and MS analysis

T cells were activated with 10ng/mL PMA and 1µM Ionomycin (Sigma-Aldrich) for 4h in the presence of 1:1000 monensin (eBioscience). Cells were stained in sterile PBS for 30min at 4**°**C with anti-CD8α (RPA-T8, BD) and Near-IR live-dead marker (ThermoFisher). Intracellular cytokine staining (ICCS) was performed with CytoFix-CytoPerm kit (BD) according to the manufacturer’s protocol. Prior to use, CytoFIX was pre-incubated on ice for 2min with 40IU/mL murine RNAse inhibitors (M0314, NEB). After 15min of incubation at 4**°**C, cells were washed once with 1xCytoPerm that was pre-incubated with 40IU/mL RNAse inhibitors. Cells were then incubated in 1xCytoPerm (pre-incubated with RNAse inhibitors) for 15 min at 4**°**C. Cells were then washed and re-suspended in Sorting buffer (2xSSC: 300mM NaCl, 30mM sodium citrate, 80IU/mL RNAse inhibitor). Antibodies directed against IL-2 (MQ1-17H12) and IFN-γ (4S.B3) were incubated overnight in Sorting buffer at 4**°**C. Cells were washed once and re-suspended in Sorting buffer prior to FACS-sorting.

To evaluate the protein recovery of fixed samples compared to fresh samples, we used 5×10^6^ PBMCs treated with CytoFix-CytoPerm kit (BD) according to the manufacturer’s protocol or left in PBS. The samples were washed once with PBS, snap-frozen in liquid nitrogen, and used for MS analysis.

### FACS-sorting

The pre-cooled FACS Aria III (BD) sorter was washed once with 70% Ethanol, followed by a 5min wash with Cleaning buffer (20xSSC (3M NaCl, 300mM sodium citrate, 400IU/mL RNAse inhibitors). Sorted cells (excluding cell doublets and dead cells) were collected in 500µL Sorting buffer. Cells were spun for 20min at 4**°**C at 4000rpm (2820g). The cell pellet was transferred in 1mL Sorting buffer in lo-Bind tubes (Eppendorf) and spun at 10.000g for 5min in table-top centrifuge (Eppendorf). Supernatant was removed, and pellet was frozen at −80**°**C until further use. Live cells were FACS-sorted according to standard procedures. Antibody staining was performed in PBS + 1% FCS for 30min at 4**°**C for live-dead marker (Invitrogen), α-CD8 (RPA-T8, SK1), α-CD29 (Mar4, BD), α-CD38 (HIT2, Biolegend), and for murine α-TCRβ when selecting for MART1-TCR expressing T cells (H57-597). Cells were washed once with culture medium and sorted on a pre-cooled FACS ARIA III sorter. Sorted cells were collected in culture medium.

### Flow-cytometry analysis

Cells were stained in PBS+1%FCS for live-dead marker and antibodies against: CD3 (UCHT1), CD8 (RPA-T8,SK1), CD29 (Mar4), CD38 (HIT2), CD55 (JS11), CD11b (ICRF44), CD27 (CLB-27), CD49a (TS2/7), CD49b (P1E6-C5), CD49d (9F10), CD39 (EbioA1), CD40L (24-31), CD45RA (HI100), CD5 (M1649), CD70 (Ki-24), CD74 (M-B741), CD80 (L307.4), CD81 (JS-81), CD103 (B-ly7), CD122 (Mik-B3), CD137 (4B4-1), CD226 (11-A8), CD360 (17A12), CCR4 (1G1), CX3CR1 (2A9-1), GITR (108-17), HLA-DR (LN3), ICOS (C398.4A), IL18R1 (H44), IL7R (HIL-7R-M21), KLRG1 (HP-3D9), lag3 (3DS223H), Ox40 (ACT35), PD-L1 (MIH1), for MART1-TCR selection murine TCR-B (H57-597). The cytokine production profile was determined by ICCS after activation with PMA-Ionomycin for 4h. Cells were prepared with CytoFIX-CytoPerm kit following manufacturer’s protocol. Cells were stained with antibodies against IFN-γ (4S.B3), IL-2 (MQ1-17H12), TNF-α (MAb11), Granulysin (DH2), Granzyme A (CB9), Granzyme B (GB11), CCL3 (CR3M), CCL4 (FL34Z3L), CCL5 (21445), Perforin (dG49). Cells were acquired on LSR II, Fortessa or Symphony (all BD) using FACS Diva v8.0.1 (BD). Data were analysed with FlowJo VX (TreeStar).

### Sample preparation for RNA-sequencing

For validation assays, cell pellets were defrosted on ice, and resuspended in proteinase K digestion buffer (20 mM Tris-HCl (pH 8.0), 1 mM CaCl2, 0.5% sodium dodecyl sulfate) and 75 µg molecular grade proteinase K (Life-Technologies). After 1h incubation at 55**°**C, Trizol-chloroform extraction was performed with 500ul according to the manufacturer’s protocol (Invitrogen). For RNA-seq analysis, RNA was isolated with the RNAeasy-FFPE kit (Quiagen) according to the manufacturer’s protocol, omitting the steps directed at deparaffinization. RNA was resuspended in RNAse-free water. RNA concentration was measured with nanodrop (ThermoFisher), and RNA integrity was determined with the RNA 6000 Nano assay on the Bioanalyser 2100 (Agilent) Sequencing was performed with stranded “FFPE-library preparation” (GenomeScan, Leiden, the Netherlands), including Ribosomal RNA depletion with NEB Next rRNA depletion kit (NEB# E6310). Sequencing was performed on Illumina HiSeq 4000, with an average depth of 21 million paired-end 150nt (PE150) reads.

### Quantitative PCR analysis

For validation experiments, cDNA preparation was performed with random hexamers using Super Script III reverse transcription (Invitrogen) according to the manufacturer’s protocol. RT-qPCR Primers were designed with the NCBI primer blast tool (Ye et al., 2012), and manufactured by Sigma. qPCR Primer pairs were used when standard curves showed an r^2^>0.98. RT-qPCR was performed with Power SYBR-green (ThermoFisher) with the standard protocol (T_m_ = 60°C for 1 min) on a 7500 Real-time qPCR system (Applied Biosystems). RT-qPCR data were analysed with 7500 Software v2.3 (Applied Biosystems). Ct difference was calculated by subtracting the Ct values from fixed or fixed-reversed to the Ct values from unfixed sample and plotted using GraphPad PRISM. Primers: *β-Actin* (Fwd: AGAGCTACGAGCTGCCTGAC, Rev.: AGCACTGTGTTGGCGTACAG) *RPS18*: (Fwd: CAGAAGTGACGCAGCCCTCTA, Rev.: AGACAACAAGCTCCGTGAAGA), *GAPDH* (Fwd: GAGACTCGTGCAATGGAGATTCT, Rev.: ACCCTGTTGCTGTAGCCA). *IL2* and *IFNG* primers were previously described (Nicolet et al., 2017).

### RNA-seq data analysis

Reads quality was inspected using fastQC version 0.11.7 (S. Andrews, https://www.bioinformatics.babraham.ac.uk/projects/fastqc/). Reads were aligned with STAR version 2.5.0a (Dobin and Gingeras, 2015) on the human genome hg38-release 92 from ENSEMBL (Zerbino et al., 2018). STAR was used to count reads per genes (option --quantMode GeneCounts) and DESeq2 (Love et al., 2014) using Wald test with p-adjusted <0.01 and log2 fold change (LFC) >0.5 to isolate differentially expressed genes. Further annotation was obtained from Ensembl BioMart (Zerbino et al., 2018; Kinsella et al., 2011) to discriminate between different types of coding and non-coding genes. Reads per million mapped reads (RPM) were calculated by dividing the read counts per gene by the total number of mapped reads.

### Mass spectrometry analysis

Sample preparation was performed as previously described (van Aalderen et al., 2017). Tryptic peptides were separated by nanoscale C18 reverse phase chromatography coupled on line to an Orbitrap Fusion Tribrid mass spectrometer (Thermo Scientific) via a nanoelectrospray ion source (Nanospray Flex Ion Source, Thermo Scientific). The MS^2^ ion count target was set to 1.5 x 10^4^. Only precursors with charge state 2–7 were sampled for MS^2^. The instrument was run in top speed mode with 3s cycles. All data were acquired with Xcalibur software.

The RAW mass spectrometry files were processed with the MaxQuant computational platform, 1.5.2.8. Proteins and peptides were identified using the Andromeda search engine by querying the human Uniprot database (downloaded February 2015). Standard settings with the additional options match between runs, Label Free Quantification (LFQ), and razor + unique peptides for quantification were selected. The generated ‘proteingroups.txt’ table was filtered for potential contaminants, reverse hits and ‘only identified by site’ using Perseus 1.5.1.6. The LFQ values were normalized using a log2 transformation. Data were imported in R and processed with the DEP package (Zhang et al., 2018), and filtered for proteins that were found in all biological replicates of at least one population. Filtered data were normalized and imputed using random draws from a Gaussian distribution centered around a minimal value (q=0.01). Differential protein expression was determined with DEP (which uses Limma) with p-adj. <0.05 and log2 fold change >0.5.

### Single cell RNA-seq analysis

Single cell RNA-seq datasets were reanalysed from Guo et al. (Guo et al., 2018) and Zheng et al (Zheng et al., 2017). Count matrix was filtered for “PTC” for peripheral blood CD8^+^ T cells, “NTC” for normal lung or liver CD8^+^ T cells. Single cell toolkit (SCTK) or ASAP (for clustering of Naïve-like cells) was used scRNA-seq data analysis (Gardeux et al., 2017; Jenkins et al., 2018). First, the batch-effect from the different patients was corrected using ComBaT in SCTK (Li et al., 2007). Cells were filtered for >1000 expressed genes per cell, keeping the 50% most expressed genes after zero removal. We then log-transformed and performed differential expression (DE) analysis (absolute Log fold change >=0.5 and p-adjusted <0.05). CD29 grouping was determined based on the “double peak” expression distribution of *ITGB1* (CD29), “PTC” from blood CD8^+^ (cut-off: 10reads/cell), “NTC” from Lung or Liver CD8^+^ cells (cut-off: 32 reads/cell (lung) and 10 reads/cell (liver)) (see Fig S7). Unsupervised clustering was performed in ASAP, using k-means (set to k=4) on tSNE. For validation, naïve-like cells were compared to the remaining “non-naïve-like” cells with ASAP (using Limma, LFC>1 and p-adjusted < 0.05). All DE genes were used for volcano-plots, and Venn diagrams show the top50 most upregulated DE genes.

### Filtering of differentially expressed gene for functional annotation

Genes encoding secreted proteins were extracted from https://www.proteinatlas.org/humanproteome/tissue/secretome (downloaded Jan. 2018), and differentially expressed genes (DEG), or proteins with an absolute(LFC)>1.5, were identified. Of note, we manually curated the list for secreted genes for obvious mis-annotations (histones, membrane, TF, RNA-binding, CD molecules, protein without protein evidence, collagen, nuclear, nucleus, ribosomal, mitochondrial). Similarly, genes and proteins encoding CD molecules were extracted with the corresponding gene and protein names (www.uniprot.org/docs/cdlis; downloaded July 2017), and DEG, or proteins with an absolute(LFC)>1.5, were identified.

Genes encoding for Transcription Factor were obtained from https://www.proteinatlas.org/humanproteome/proteinclasses (downloaded Jan. 2019), and filtered from the DEG. LncRNA were filtered from DEG, excluding protein-coding genes and TCR/IG pseudogenes. Genes of experimentally validated RNA-binding proteins were obtained from (Castello et al., 2016; Perez-Perri et al., 2018; Gerstberger et al., 2014), and filtered from the DEG.

### Survival analysis

Oncolink (Anaya, 2016) was used to assess the effect of *CD8B* expression on different types of cancer. A Cox regression analysis for CD8B was perfomed and the –log(FDR) was used to identify responsive cancer types (FDR<0.01).

Datasets for SKCM (skin cutaneous melanoma) RNA-sequencing and patient survival were obtained from TCGA (www.cancer.gov/tcga) using UCSC Xena (xenabrowser.net/datapages/; BioRxiv: doi: 10.1101/326470v5). Gene signature of CD29+ CD8+ T cells (Figure 5G) was used to isolate the gene expression from the RNA-sequencing SKMC TCGA dataset (in TPM+1). TPM+1 counts were transformed into a gene-wise Z-score. The average of all Z-score was calculated per patient. Patients were then stratified in two groups (high or low CD29 signature expression) using the median of the Z-score average (as described in (Guo et al., 2018)). Patients were also stratified for CD8^+^ T cell infiltrates using estimates obtained by CIBERSORT (Newman et al., 2015) on SKCM TPM+1 counts. Survminer R package was used to prepare and visualize the patient survival with Kaplan-Meier plot. Log-rank test was used to determine statistical differences in survival between group.

### Generation of MART1-TCR expressing T cells

PBMCs from individual donors were activated for 48h with αCD3/αCD28 as described above. Cells were harvested and retrovirally transduced with the MART1-TCR, as previously described (Gomez-Eerland et al., 2014). Briefly, non-tissue cultured treated 24 well plates were coated with 50μg/mL Retronectin (Takara) overnight and washed once with 1mL/well PBS. 300µL/well viral supernatant was added to the plate, and plates were centrifuged for 1h at 4**°**C at 4000 rpm (2820g). 0.5×10^6^ T cells were added per well, spun for 10min at 1000rpm, and incubated overnight at 37**°**C. The following day, cells were harvested and cultured in T25 flasks at a concentration of 0.8×10^6^ cells/mL for 6-8 days in presence of 10ng/mL rhIL-15.

### Functional assays with MART1-TCR expressing T cells

MART1-TCR transduced CD8^+^ T cells were FACS-sorted based on TCR expression, and rested over-night in medium at 37**°**C. Cytokine production was determined, as described above, by ICCS after 6h of co-culture with HLA-A2^+^ MART1^hi^ Mel 526 (MART1+), or HLA-A2^-^ MART1^lo^ Mel 888 (MART1-) tumor cell lines (Marincola et al., 1994), in a 3 to 1 effector to target (E:T) ratio. For CD107α staining, co-cultured cells were supplemented with anti-CD107a, Brefeldin A and Monensin, in the culture medium for 6h. Cells were subsequently measured by flow cytometry. For killing assays, total CD8^+^ MART1 TCR-engineered T cells were FACS-sorted or sorted based on their CD29 and CD38 expression profile (CD29^+^ and CD38^+^). Tumor cells were labelled with 1.5μM CFSE for 10 min at 37**°**C in FCS-free medium and washed 3 times with warm culture medium. 15×10^3^ tumor cells were co-cultured with MART1-TCR^+^ FACS-sorted T cells for 20h, in a 3:1, 1:1 and 0.3:1 E:T ratio. Dead cells were quantified by flow-cytometry with Near IR live-dead marker on CFSE^+^ tumor cells.

### Statistical analysis and Data visualisation

Data generated with flow cytometry were analysed with paired t-test, repeated measurement ANOVA using GraphPad PRISM version 7. Differences were considered significant with a p-value < 0.05.

Plots were generated with ggplot2 version 3.0, DEP version 1.2.0, and with GraphPad. Heatmaps were generated with pHeatmap version 1.0.10 and DEP. Venn diagrams were generated with (http://bioinformatics.psb.ugent.be/webtools/Venn/).

## SUPPLEMENTARY MATERIALS

**Fig. S1**: RNA recovery after fixation of T cells.

**Fig. S2**: Differential cytokine production and surface molecule expression profile of IL2 and IFNG producers

**Fig. S3**: Characterization of CD29^+^ and CD38^+^ T cells

**Fig. S4**: Ex vivo analysis of CD29^+^CD8^+^ T cells.

**Fig. S5**: CX3CR1 marks a subset of CD29^+^CD8^+^ T cells

**Fig. S6**: CD29^+^MART1-specific T cells are superior cytokine producers and show higher killing capacity than CD38^+^ T cells

**Table S1**: Differentially expressed genes and proteins between DN, IL2, IFNG, and DP populations.

**Table S2**: List of differentially expressed genes and proteins annotated as: secreted proteins, transcription factors and RNA-binding proteins.

**Table S3**: Differentially expressed genes between ITGB1high and low from scRNA-seq data.

## ACKNOWLEDGMENTS

We would like to thank the blood donors for donation and E.Mul, M.Hoogenboezem and S.Tol for FACS sorting. We are grateful to B. Popovic and D. Amsen for critical reading of this manuscript.

## AUTHOR CONTRIBUTIONS

B.P.N. and M.C.W. designed experiments; B.P.N., A.G. F.P.J.v.A. and M.v.d.B. performed experiments and analysed data, R.G.-E. and T.N.M.S. provided the MART1 TCR system. M.C.W. directed the study; B.P.N. and M.C.W wrote the manuscript.

## DATA AVAILABILITY

All sequencing data are deposited on NCBI GEO under the accession no: GSE125497, and all MS data were deposited on PRIDE PXD012874.

