## Supplementary material for "CD29 marks superior cytotoxic human CD8^+^ T cells": Fig. S1-6

### Nicolet et al., Figure S1

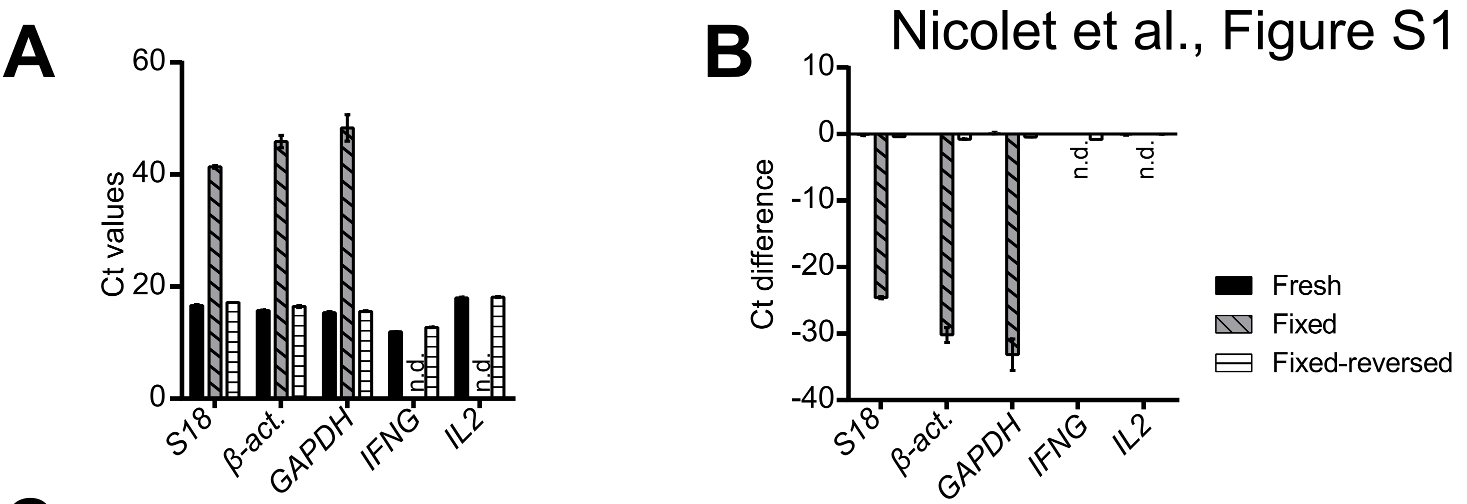


C

| | | RNA QC | | | | | Library QC | | FastQc | Seq. Depth | Alignment |
|---|---|---|---|---|---|---|---|---|---|---|---|
| | | Recovered RNA (ng) | 260/280 | 260/230 | RIN (QC in house) | RIN (QC by company) | Peak (nt) | Avg. size(nt) | All within norms? | Million reads | STAR (% mapped) |
| 001-001 | IL2-1 | 1590,4 | 1,92 | 2,32 | 3,1 | 3,3 | 402 | 414 | Yes | 20,8 | 93,61 |
| 001-002 | IFNG-1 | 1383,2 | 1,96 | 2,16 | 2,6 | 3,8 | 404 | 421 | Yes | 23,7 | 93,54 |
| 001-003 | DP-1 | 3137,4 | 2,03 | 2,42 | 3,2 | 3,8 | 416 | 423 | Yes | 15,7 | 92,32 |
| 001-004 | DN-1 | 1451,8 | 1,98 | 2,68 | 2,8 | 3,8 | 416 | 426 | Yes | 24,5 | 93,42 |
| 001-005 | IL2-2 | 2748,2 | 2,01 | 2,32 | 7,5 | 7,3 | 415 | 415 | Yes | 22,4 | 91,32 |
| 001-006 | IFNG-2 | 1604,4 | 1,98 | 2,69 | 7,2 | 8,2 | 417 | 422 | Yes | 17,7 | 91,86 |
| 001-007 | DP-2 | 5090,4 | 2,04 | 2,22 | 6,7 | 7,7 | 417 | 420 | Yes | 25,5 | 90,71 |
| 001-008 | DN-2 | 1577,8 | 1,99 | 2,57 | 7,3 | 8 | 416 | 424 | Yes | 28,5 | 91,99 |
| 001-009 | IL2-3 | 2031,4 | 1,91 | 2,61 | N.A | 5,5 | 416 | 420 | Yes | 18,2 | 93,87 |
| 001-010 | IFNG-3 | 1650,6 | 1,96 | 2,55 | 6,4 | 7,7 | 416 | 424 | Yes | 18,3 | 91,36 |
| 001-011 | DP-3 | 5278 | 1,99 | 2,47 | 6,4 | 7 | 403 | 412 | Yes | 17,4 | 91,50 |
| 001-012 | DN-3 | 753,2 | 1,92 | 2,07 | 6,6 | 7,7 | 400 | 428 | Yes | 24,3 | 92,25 |

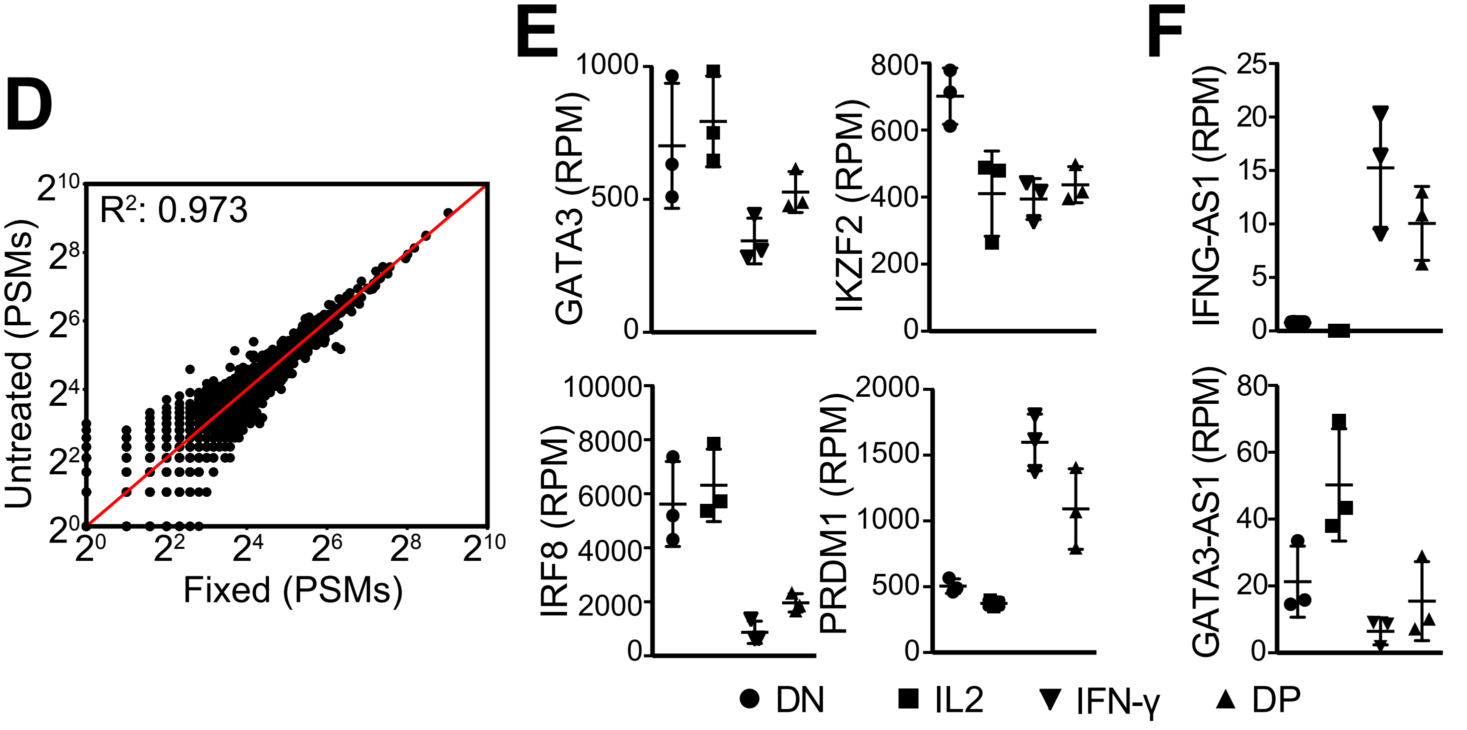

### Nicolet et al., Figure S2

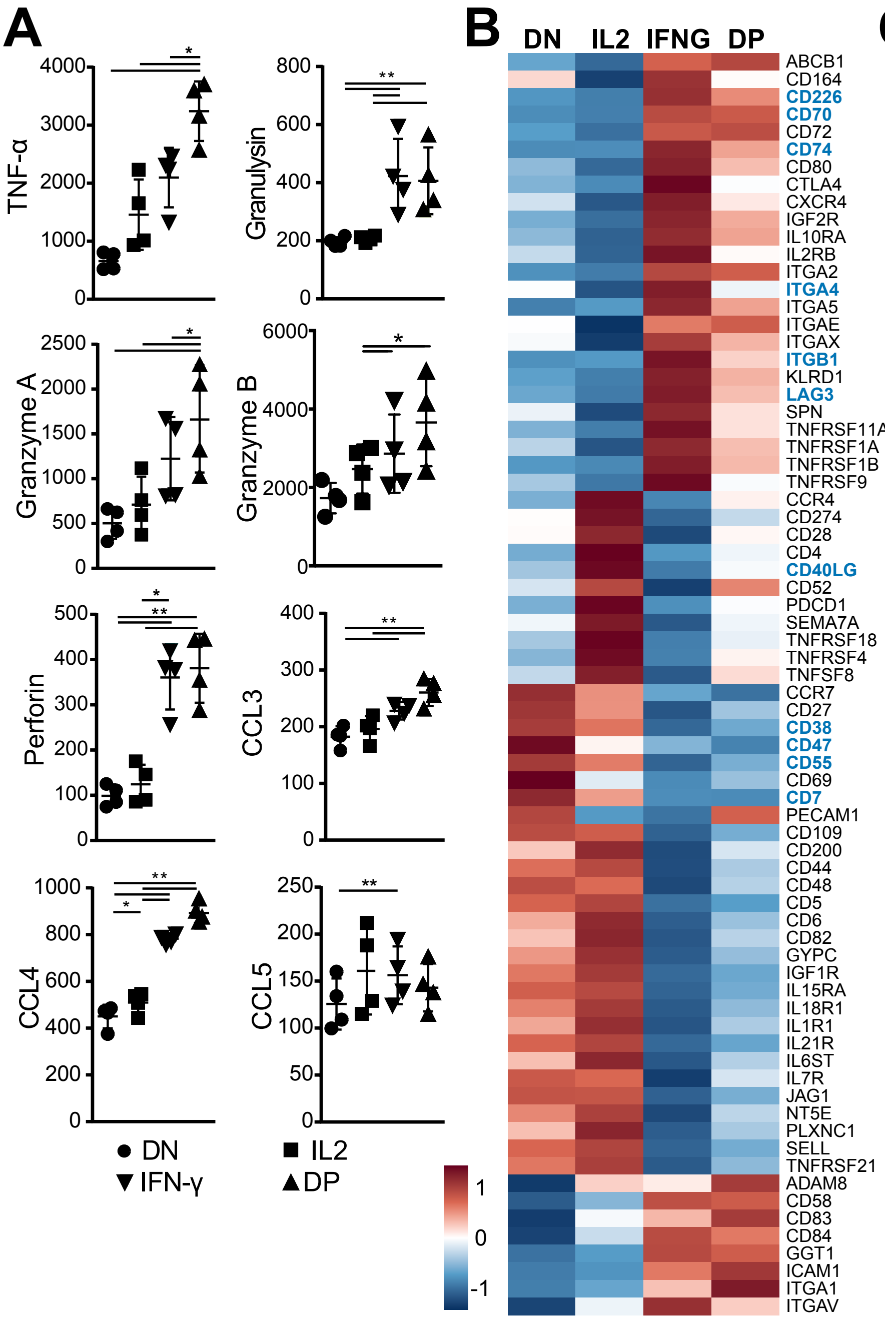


C

| Marker | In MS | In RNA-seq | Staining | In line with expression data | Selection of IFNg/IL2 production? | Clone |
|---|---|---|---|---|---|---|
| **CD29 (ITGB1)** | yes | yes | Good | Yes | Good | mar4 |
| **CD38** | yes | yes | Good | Yes | Good | HIT2 |
| CD39 | Yes | No | Good | Yes | No | EbioA1 |
| CD40LG | yes | yes | Weak | Yes | No | 24-31 |
| CD49d | yes | yes | Good | Yes | Weak | 9F10 |
| CD55 | yes | yes | Good | Yes | Good | JS11 |
| CD70 | yes | yes | Good | Yes | No | Ki-24 |
| CD74 | yes | yes | Good | Yes | Good | M-B741 |
| CD81 | Yes | no | Weak | Yes | Weak | JS-81 |
| CD226 | yes | yes | Bad | NA | NA | 11A8 |
| HLA-DR | yes | yes | Good | Yes | Medium | LN3 |
| Lag3 | yes | yes | Bad | NA | NA | 3DS223H |
| CCR4 | no | yes | Good | Yes | Weak | 1G1 |
| CD103 | no | yes | Good | Yes | Weak | B-ly7 |
| CD11b (ITGAM) | no | yes | Good | Yes | Weak | ICRF44 |
| CD122 (IL2RB) | no | yes | Good | Yes | Weak | Mik-B3 |
| CD127 (IL7R) | no | yes | Bad | NA | NA | HIL-7R-M21 |
| CD134 (Ox40) | no | yes | Good | Yes | No | ACT35 |
| CD137 (4-1BB) | no | yes | Weak | Yes | No | 4B4-1 |
| CD218a (IL18R1) | no | yes | Bad | NA | NA | H44 |
| CD274 (PD-L1) | no | yes | Weak | Yes | No | MIH1 |
| CD357 (GITR) | no | yes | Bad | NA | NA | 108-17 |
| CD360 (IL21R) | no | yes | Bad | NA | NA | 17A12 |
| CD49a | no | yes | Good | Yes | Weak | TS2/7 |
| CD49b | no | yes | Good | Yes | Weak | P1E6-C5 |
| CD5 | no | yes | Weak | Yes | No | M1649 |
| CD80 | no | yes | Weak | Yes | No | L307.4 |
| CD94 (KLRD1) | no | yes | Good | Yes | No | HP-3D9 |

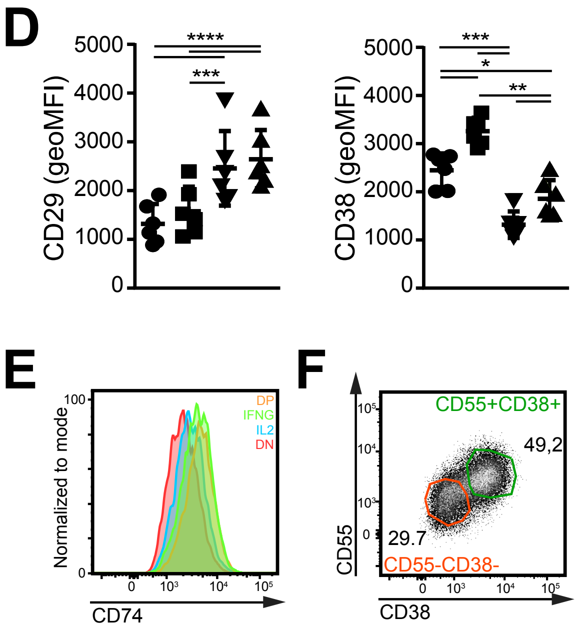

### Nicolet et al., Figure S3

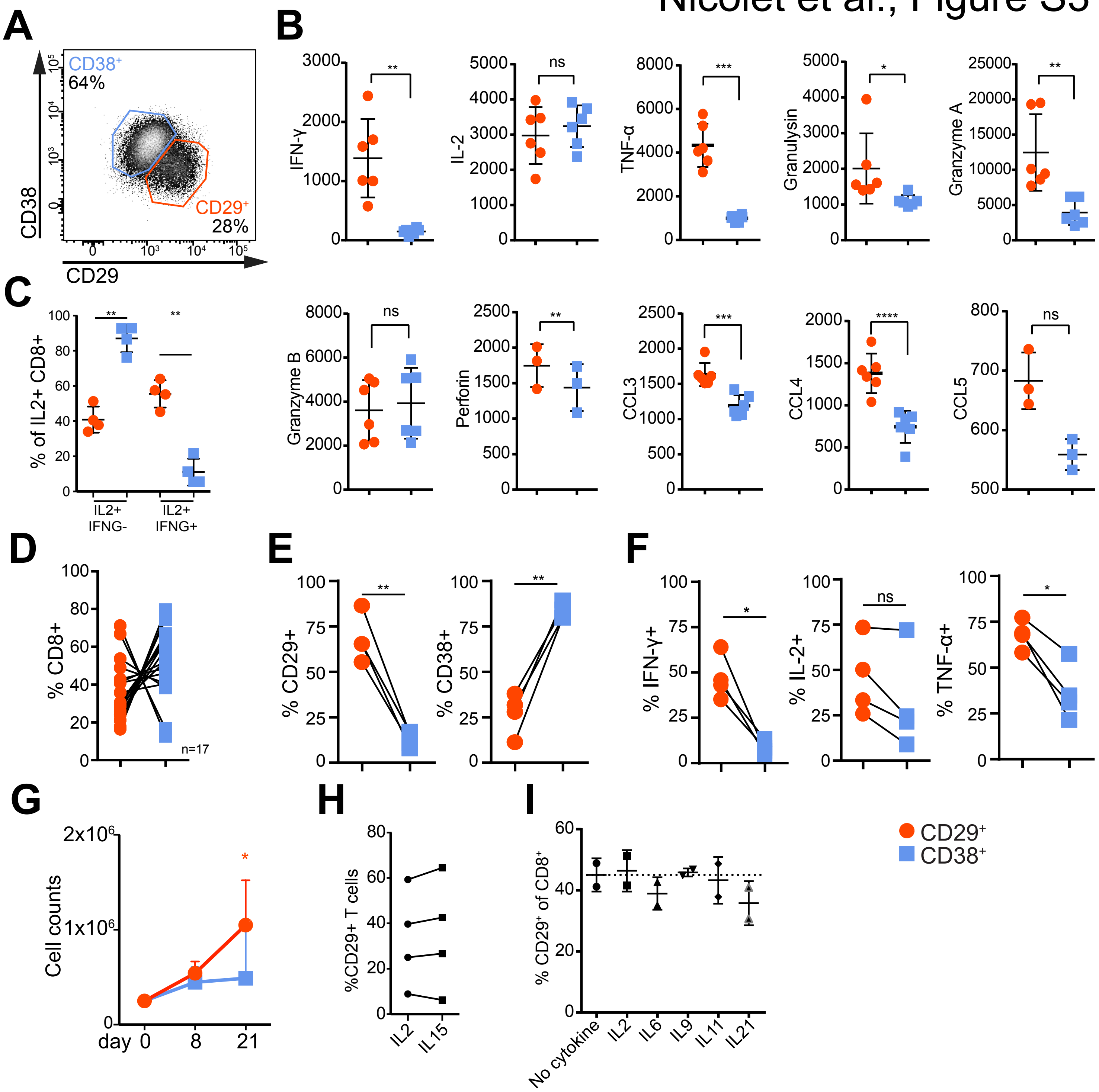

### Nicolet et al., Figure S4

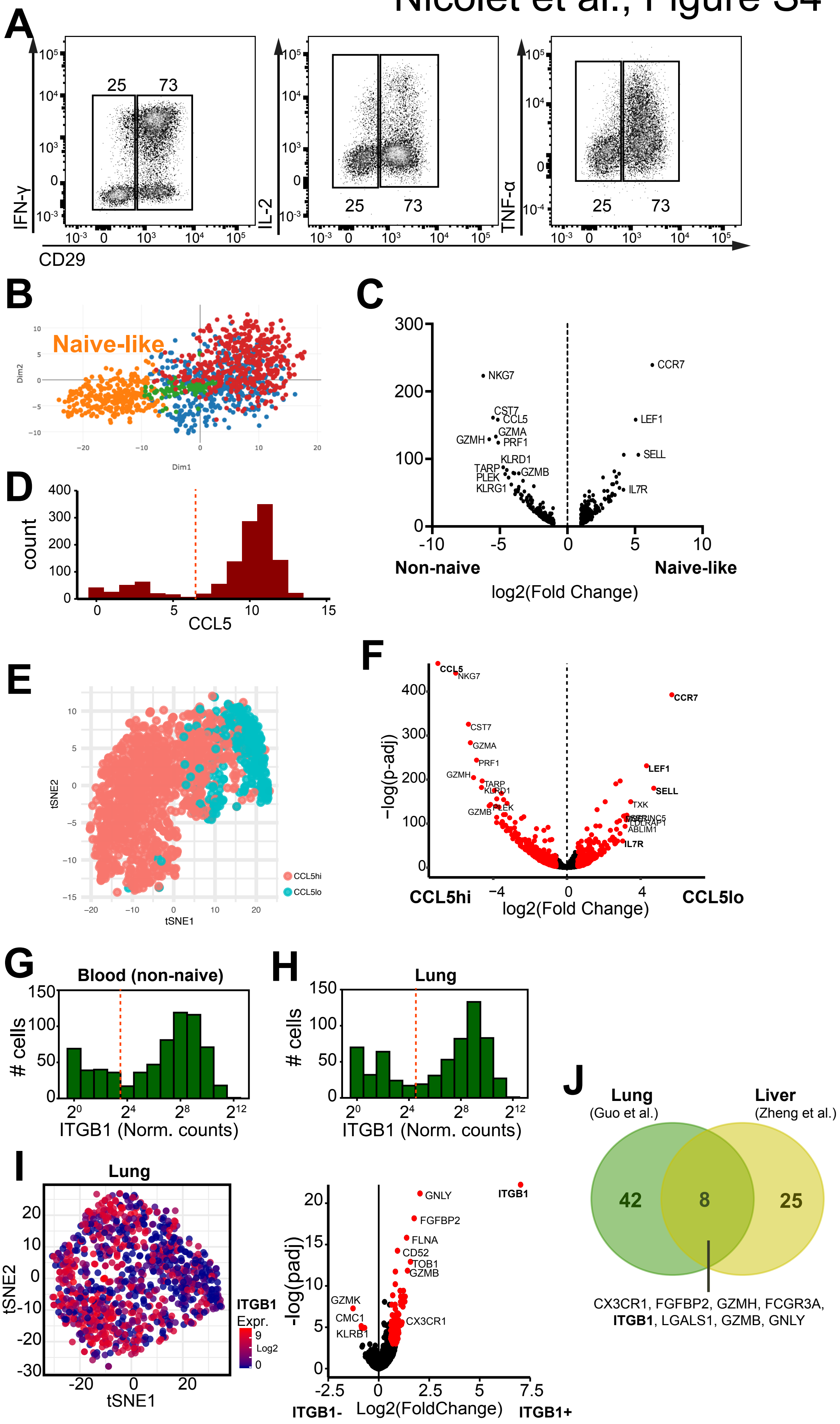

### Nicolet et al., Figure S5

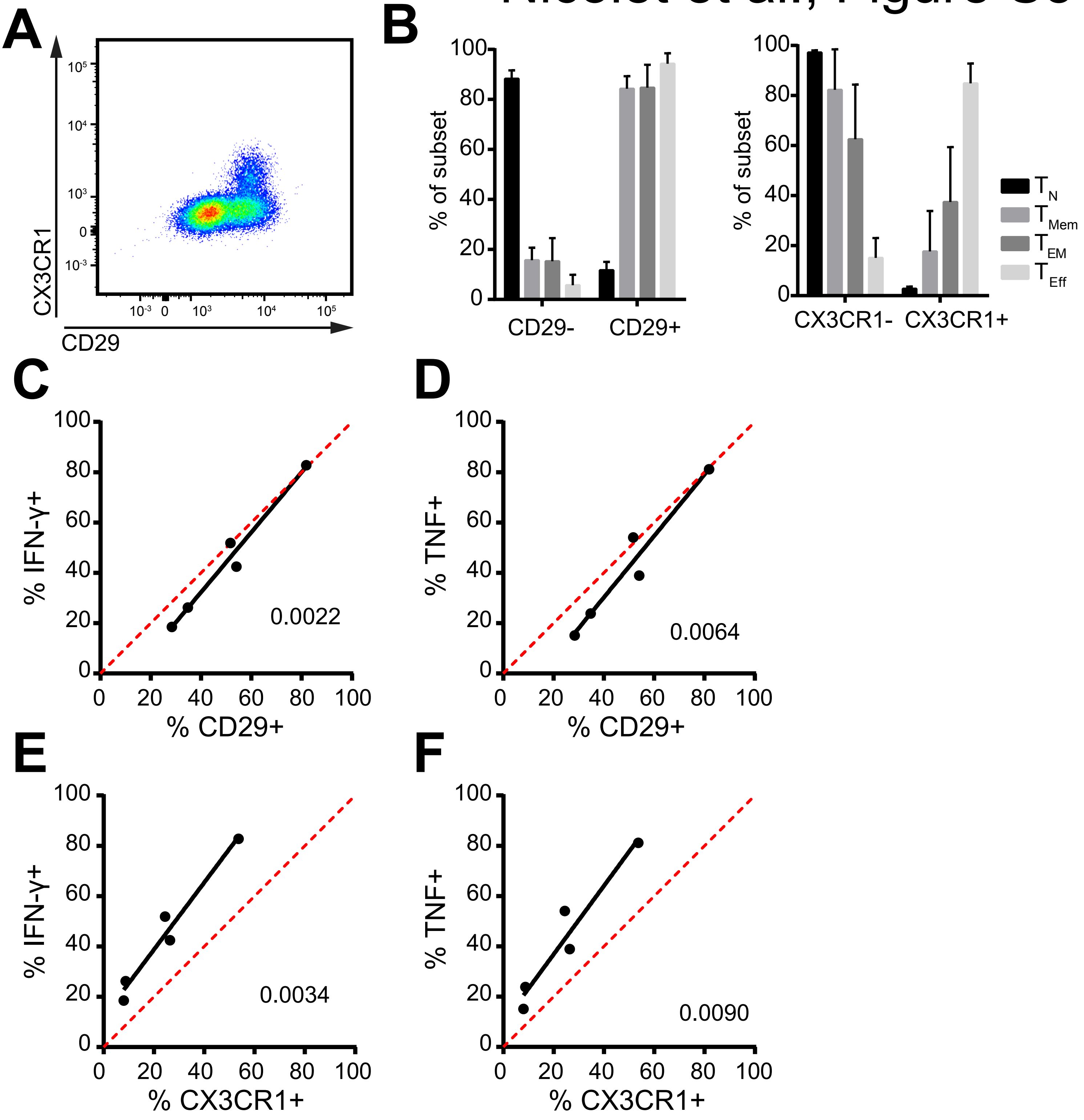

### Nicolet et al., Figure S6

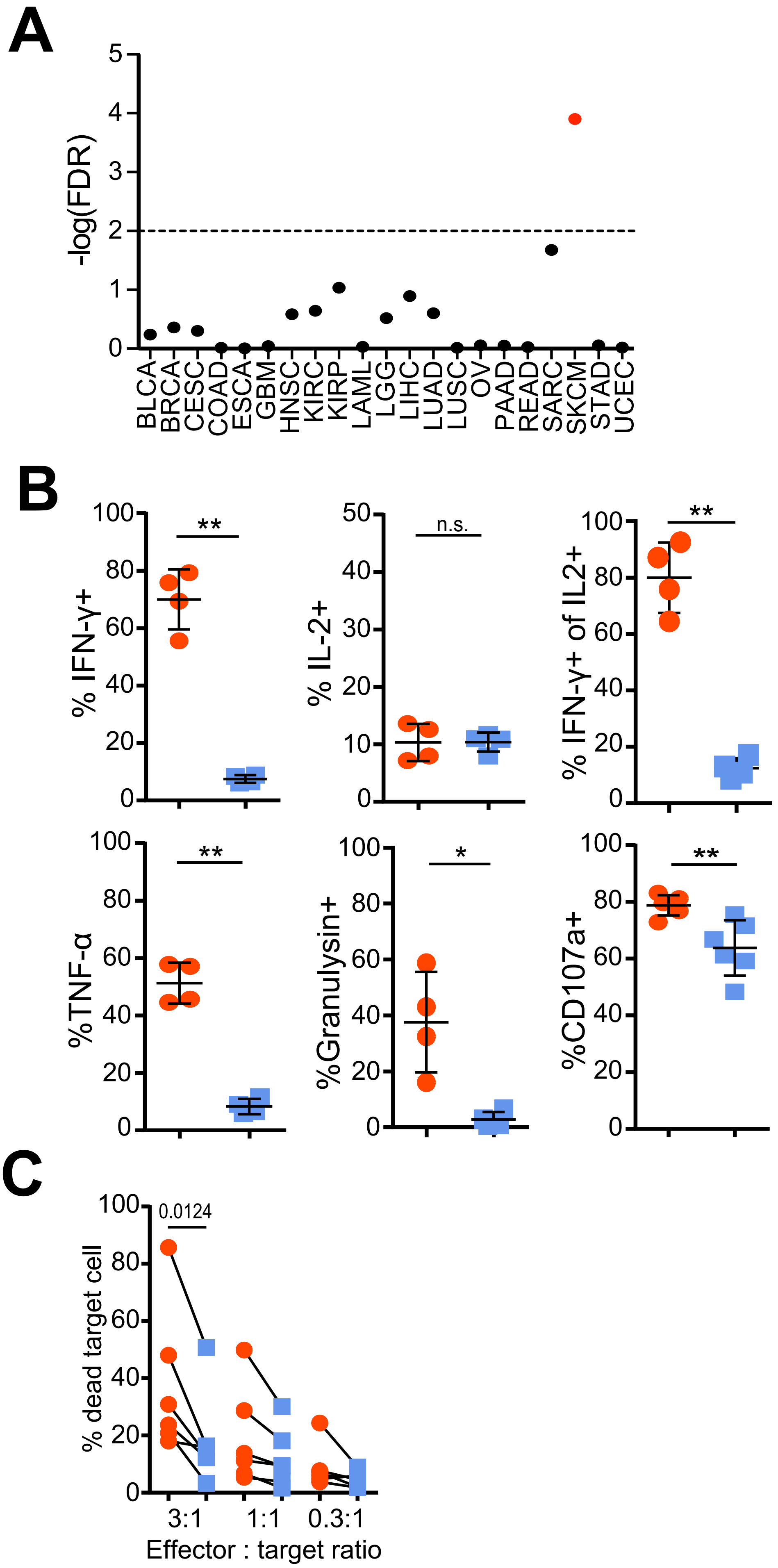
